# Metabolic control of drug resistance by a mycobacterial ion channel

**DOI:** 10.64898/2026.03.10.710584

**Authors:** Alexandre Gouzy, Shuqi Li, James Chen, Allen Na, Anas Saleh, Zachary A. Azadian, Kayan Tam, Vanisha Munsamy-Govender, Nicholas C. Poulton, Michael A. DeJesus, Dirk Schnappinger, Kyu Y. Rhee, Sabine Ehrt, Jeremy M. Rock

**Affiliations:** Department of Microbiology and Immunology, Weill Cornell Medical College, New York, NY, 10065, USA; Laboratory of Host-Pathogen Biology, The Rockefeller University, New York, NY, 10065, USA

## Abstract

Pyrazinamide (PZA) is a cornerstone of modern tuberculosis therapy, yet its context-dependent activity has obscured both its mode of action and resistance mechanisms. Using a host-mimicking culture system integrated with genome-wide CRISPRi profiling, metabolomics, and comparative genomics, we identify a previously unrecognized driver of PZA resistance in humans: loss of the ion channel Rv2571c. Rv2571c mediates α-ketoglutarate efflux, amplifying PZA-induced cytoplasmic acidification under host-relevant acidic conditions. Loss-of-function mutations confer resistance in vitro and in vivo and are under positive selection in clinical isolates, establishing this pathway as a resistance determinant in patients. Together, these findings define a novel, ion channel–mediated resistance mechanism, establish cytoplasmic acidification as the basis of PZA killing, and inform resistance detection and treatment-shortening drug development.

## INTRODUCTION

Tuberculosis (TB) remains the leading cause of death from an infectious disease, and the continued emergence of antibiotic resistance threatens global control efforts^1^. TB is caused by the bacterial pathogen *Mycobacterium tuberculosis* (Mtb) and is treated with a prolonged multidrug regimen comprising rifampicin, isoniazid, ethambutol, and pyrazinamide (PZA). Among these agents, PZA plays a uniquely important role by shortening treatment duration from nine to six months and reducing relapse^2–4^, potentially by targeting non-replicating, drug-tolerant Mtb^5–8^ and accumulating within granulomas and phagosomes of infected macrophages^9,10^. Despite decades of clinical use, PZA’s mechanism of action remains controversial, and physiologically relevant mechanisms of resistance are incompletely defined^11–13^.

PZA is a prodrug that is converted by the mycobacterial nicotinamidase/pyrazinamidase PncA into its active form, pyrazinoic acid (POA)^14^. Loss-of-function mutations in *pncA* account for the majority of clinically observed PZA resistance (PZA-R)^15–17^. However, *pncA* is dispensable for bacterial fitness, resulting in a diffuse mutational landscape^16,18^ that complicates the development of targeted molecular diagnostics analogous to those used to detect rifampicin-resistance^19^. Compounding this challenge, not all *pncA* polymorphisms confer PZA resistance^16,18^, and 10-30% of phenotypically PZA-resistant clinical isolates lack *pncA* mutations^17,20^, indicating that additional resistance mechanisms remain to be identified. Together, these features have limited the routine molecular detection of PZA resistance in clinical practice^21–24^.

Beyond resistance detection, the mechanism of action of PZA remains controversial^11–13^. Since its introduction in the 1950s, proposed models include inhibition of trans-translation^25^, disruption of fatty acid^26^ or coenzyme A synthesis^27–29^, and cytoplasmic acidification^8,30^. This lack of consensus is driven, at least in part, by the experimental challenges inherent to studying PZA. PZA is largely inactive in standard laboratory media buffered near neutral pH^8,30–34^, despite its potent activity in animal models and patients^3,10,35^. In contrast, acidic conditions strongly enhance PZA activity *in vitro*^8,30,32,33^, consistent with the acidic environments encountered by Mtb within macrophage phagosomes (pH ∼4.5–6.2) during infection^36–38^. Inhibition of phagosomal acidification correspondingly reduces PZA efficacy, underscoring the importance of pH in drug action^9,39^.

To address these challenges, we built on our prior work^40^ showing that host-derived lipids sustain Mtb growth under acidic conditions and developed a more physiologically relevant culture system that enables robust and reproducible measurement of PZA activity *in vitro*. Using this system, we validated acidic pH as a strict requirement for PZA-mediated killing. Leveraging this platform in combination with genome-wide CRISPR interference (CRISPRi) chemical–genetic profiling, comparative genomics, and metabolomics, we identify the previously uncharacterized ion channel Rv2571c as a clinically relevant determinant of PZA susceptibility. We show that Rv2571c facilitates a net efflux of α-ketoglutarate, a central TCA cycle metabolite, thereby sensitizing Mtb to PZA-induced cytoplasmic acidification under acidic conditions. Loss of Rv2571c function confers PZA resistance *in vitro*, during macrophage infection, and in mice, while gain of function results in hypersusceptibility. Together, these findings reveal a metabolite-driven mechanism linking ion channel activity, central carbon metabolism, and cytoplasmic pH homeostasis to antibiotic susceptibility, with implications for improving resistance detection and guiding the development of PZA-like therapeutics that shorten treatment.

## RESULTS

### Optimizing *in vitro* measurement of PZA activity and resistance

Accurate phenotypic assessment of PZA activity has long been limited by the drug’s strong dependence on environmental conditions. Current clinical assays typically rely on media acidified to approximately pH 5.9^41,42^. However, these tests are poorly reproducible and prone to false-positive resistance calls, likely due to medium alkalinization and the consequent loss of PZA activity^21,22,24^. More acidic conditions might preserve PZA efficacy but prevent Mtb growth in standard laboratory media that rely on glycerol and glucose as primary carbon sources^40,43–47^.

We previously showed that host-relevant lipids enable Mtb growth under acidic conditions^40^. Building on this work, we asked whether lipid-supplemented media could support both bacterial growth and robust PZA activity *in vitro*. Consistent with prior reports^8,30,32,33^, PZA exhibited no detectable activity at neutral pH, regardless of the carbon source (**Fig. 1A,B**). At pH 5.0, Mtb failed to grow in glycerol-containing media, and PZA showed only limited bactericidal activity. In contrast, supplementation with the host relevant lipid oleic acid (OA, 200 µM, replenished every 2–3 days) supported robust Mtb growth at pH 5.0 and led to potent PZA-mediated killing, resulting in complete sterilization of cultures (**Fig. 1A,B**). Replacing OA with alternative host-relevant lipids, including palmitic acid or cholesterol, yielded comparable PZA activity in acidic media (**Fig. S1A**). We next examined the relationship between pH and PZA activity in OA-containing media. Both growth inhibition and killing increased progressively as pH decreased, confirming a strict pH dependence of PZA efficacy (**Fig. S1B,C**). These data demonstrate that growth with host-relevant lipids enables reliable measurement of PZA activity under acidic conditions.

**Fig. 1.**
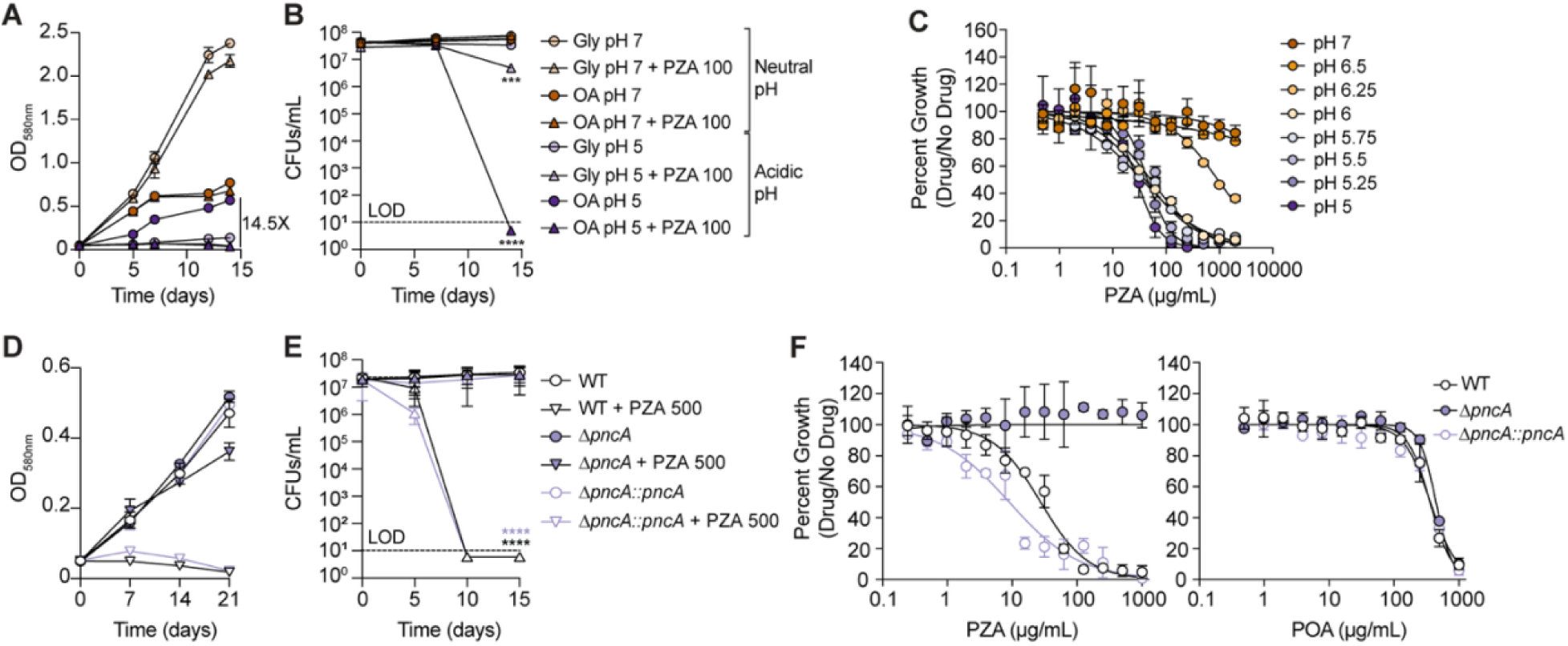
A robust assay to detect pyrazinamide (PZA) activity. (A-B) Growth (A) and survival (B) of wild-type (WT) Mtb in 7H9 medium supplemented with glycerol (Gly, ∼22mM) or oleic acid (OA, 200µM) at either pH 7 or pH 5 and in the presence or absence of PZA (100 μg/mL). (C) Growth of Mtb across a range of pH values and PZA concentrations in 7H9 medium supplemented with 3 mM OA and 50 g/L of BSA (HBO medium). (D-E) Growth (D) and survival (E) of WT Mtb, *pncA* knockout (Δ*pncA*), and the complemented strain (Δ*pncA*::*pncA*) in 7H9 medium supplemented with oleic acid (OA, 200µM) at pH 5, in the presence or absence of PZA at 500 μg/mL. (F) Dose-response curves of indicated Mtb strains treated with PZA or pyrazinoic acid (POA) in HBO medium at pH 5. In panels A, B, D, and E, 200 μM OA was replenished every 2-3 days. Panels A, B, D, and E are means ± s.d of three independent experiments. Panels C and F are means ± s.d of an experiment performed in triplicate and representative of three independent experiments. In panels B and E, statistical significance was assessed by one-way analysis of variance (ANOVA) followed by Tukey’s post-hoc test. ***, adj-*P* < 0.0005, ****, adj-*P* < 0.0001. LOD: Limit of detection.

Repeated OA supplementation is not readily compatible with small-volume or high-throughput assays. To address this limitation, we increased bovine serum albumin (BSA) concentrations tenfold, leveraging albumin’s capacity to buffer fatty acid toxicity^48,49^ while approximating physiological BSA plasma levels^50,51^. Under these conditions, Mtb grew robustly in the presence of millimolar OA at both neutral and acidic pH (**Fig. S1D**). We refer to this formulation as High-BSA-Oleate (HBO) medium.

Despite albumin’s drug-binding properties^52^, increasing BSA concentration did not significantly alter the PZA minimum inhibitory concentration (MIC_90_; **Fig. S2A**), although bactericidal activity was modestly reduced, consistent with sequestration of free OA, which can be toxic at high levels (**Fig. S2B,C**). PZA-mediated killing remained stronger in OA-containing media than in carbon-free conditions, indicating that OA potentiates but is not strictly required for PZA activity (**Fig. S 2C**). Using HBO medium, we determined the pH threshold for PZA efficacy and found that PZA activity declined sharply above pH 6.25 and was abolished at pH ≥ 6.50 (**Fig. 1C**; **Table S1**), consistent with its poor activity in standard laboratory media (7H9 medium ∼ pH 6.6).

To disentangle the role of bacterial replication from environmental pH, we assessed PZA activity in a non-replicating Mtb model. Under these conditions, acidic pH alone was sufficient to confer PZA bactericidal activity, demonstrating that active bacterial growth is not required for killing (**Fig. S2D**).

Finally, we evaluated whether these lipid-based acidic media enable reliable detection of PZA resistance. Using both OA-supplemented and HBO media at low pH, we found that a Δ*pncA* mutant was fully resistant to PZA yet remained susceptible to pyrazinoic acid (POA), whereas complementation restored PZA sensitivity (**Fig. 1D-F**). These results confirm that PZA activity in this system remains strictly dependent on PncA-mediated drug activation and demonstrate the feasibility of robust phenotypic detection of PZA resistance *in vitro*.

### Rv2571c is a clinically relevant determinant of pyrazinamide susceptibility

To identify bacterial determinants of PZA susceptibility under more physiologically relevant conditions, we combined our lipid-rich, acidic growth model with a genome-wide CRISPRi screening platform. This CRISPRi library consists of 96,700 unique single guide RNAs (sgRNAs) targeting approximately 98% of all annotated Mtb genes^53,54^. Following target pre-depletion, the pooled CRISPRi library was exposed to a subinhibitory concentration of PZA (50 µg/mL) in OA-containing acidic medium, followed by an outgrowth phase in standard media without PZA to maximize detection of resistant or sensitized strains (**Fig. 2A**). This screen identified 80 genes for which knockdown increased PZA susceptibility and 13 genes for which knockdown conferred resistance (**Fig. 2B**; **Data S1**). As expected, the top resistance-conferring hit was *pncA* (**Fig. 2B**). Among sensitizing hits, we validated several candidates, including *pckA*, *hupB*, and *blaR*, using deletion mutants (**Fig. S3**), although the mechanisms underlying their effects on PZA susceptibility remain to be defined. Together, these results validate the performance of the CRISPRi screen for identifying genetic determinants of PZA susceptibility.

**Fig. 2.**
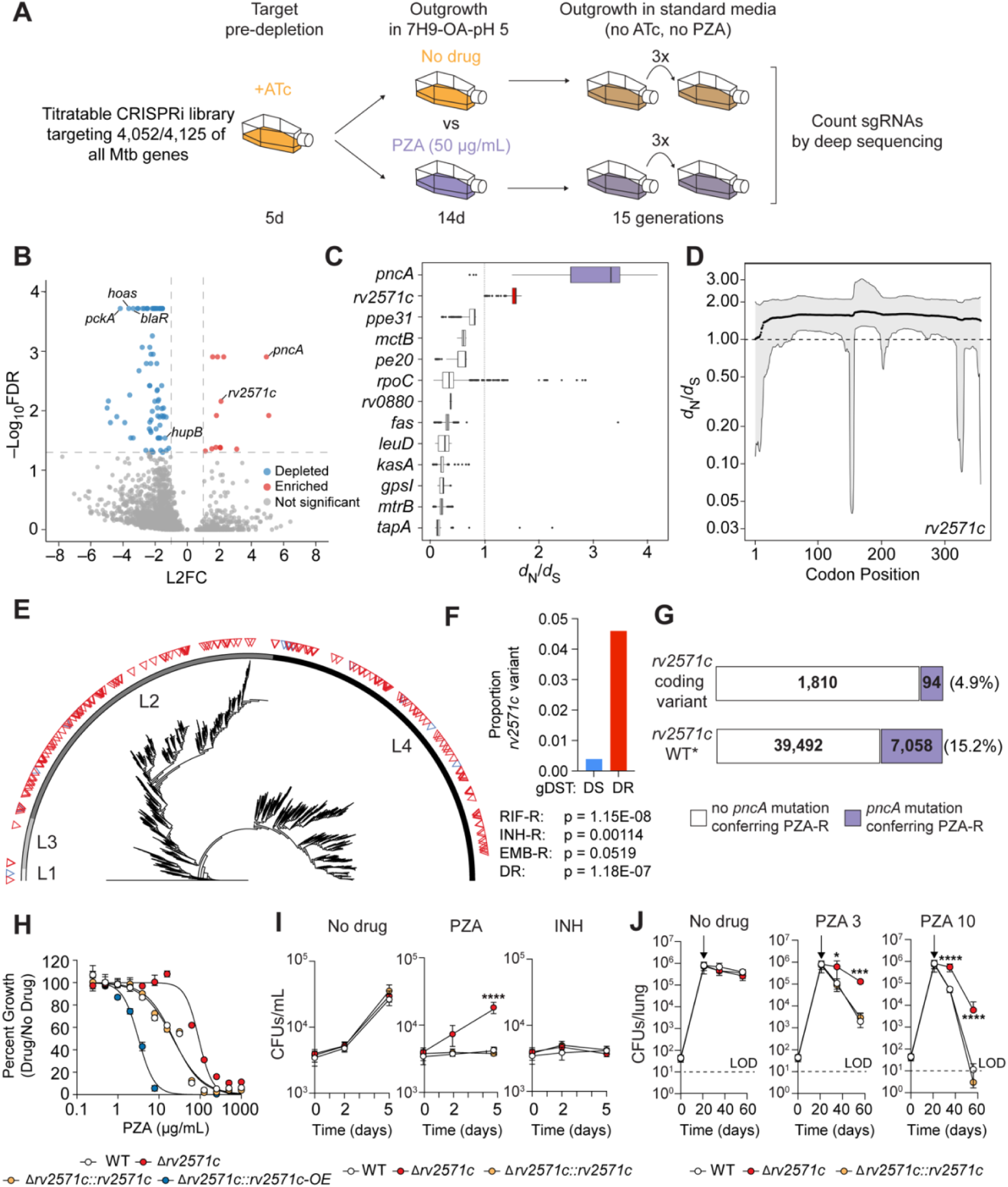
Rv2571c is a clinically relevant determinant of PZA susceptibility. (A) Schematic of the Mtb CRISPRi chemical-genetic screen to identify bacterial determinants of PZA activity. An anhydrotetracycline (ATc)-inducible, genome-wide CRISPRi library was used to systematically and tunably repress nearly all Mtb genes. ATc was added for 5 days to pre-deplete target gene products prior to drug exposure. Cultures were then split and treated with either DMSO or PZA (50 μg/mL) in acidic 7H9 medium containing OA for 14 days in the continued presence of ATc. Cells were subsequently washed and outgrown for ∼15 generations in standard 7H9 medium lacking ATc and PZA. Genomic DNA was harvested, and sgRNA abundances were quantified by deep sequencing. Chemical-genetic interactions were identified using MAGeCK. (B) Volcano plot showing log2 fold change (L2FC) values and false discovery rates (FDR) for each Mtb gene comparing PZA treatment to the no drug condition. (C) Ratio of nonsynonymous (dN) to synonymous (dS) substitutions for genes whose knockdown conferred relative PZA resistance in the CRISPRi screen. dN/dS values were calculated using GenomegaMap on a collection of 49,481 clinical Mtb genomes. (D) Codon-level dN/dS substitution rates across the *rv2571c* coding sequence. (E) Phylogenetic tree of 5,413 clinical Mtb isolates. Drug resistance was inferred based on the presence of mutations known to confer resistance to rifampicin (RIF), PZA, isoniazid (INH), ethambutol (EMB), ofloxacin, streptomycin, kanamycin, or capreomycin. Red triangles denote drug-resistant isolates harboring a single-nucleotide polymorphism (SNP) or indel in *rv2571c*, whereas blue triangles denote drug-susceptible isolates with *rv2571c* variants. L = Mtb lineage. (F) Frequencies and association of *rv2571c* variants in drug-sensitive (DS) and drug-resistant (DR) Mtb clinical isolates. Approximately 4.5% of all DR Mtb isolates harbor variants in *rv2571c*. Statistical associations of *rv2571c* variants with resistance (−R) to specific drugs were assessed using PySeer. (G) Frequency of predicted loss-of-function *pncA* mutations in clinical Mtb isolates stratified by *rv2571c* genotype (wild type versus coding variant). Isolates harboring an *rv2571c* variant are ∼3.4-fold less likely to carry a *pncA* mutation than *rv2571c* wild-type isolates (Fisher’s exact test, OR = 0.29, p = 5.18 × 10⁻⁴⁴). (H) Dose-response curves for the indicated Mtb strains treated with PZA in HBO pH 5.5 medium. Values are from experiments performed in triplicate and are representative of three independent experiments. (I) Growth of indicated Mtb strains in murine bone marrow-derived macrophages treated with DMSO, PZA, or INH. Statistical significance was assessed by one-way ANOVA followed by Tukey’s post-hoc test. ****, adj-*P* < 0.0001. Values are from experiments performed in triplicate and are representative of three independent experiments. (J) C57BL/6 mice were infected by aerosol with the indicated Mtb strains and bacterial burdens were quantified from lung homogenates at days 21, 35, and 56 post-infection. Mice were either left untreated (No drug) or treated with PZA administered in the drinking water at 3 mg/mL (middle) or 10 mg/mL (right), beginning at day 21 post-infection (arrows). Dashed lines denote the limit of detection (LOD). Statistical significance was assessed by one-way ANOVA followed by Tukey’s post-hoc test. *, adj-*P* < 0.05, **, adj-*P* < 0.005, ***, adj-*P* < 0.0005, ****, adj-*P* < 0.0001. Data represent mean ± s.d. from four mice per strain per time point and are representative of two independent experiments. In panels E, F, and G, a lineage 3–specific *rv2571c* SNP F163S, which is fixed across lineage 3 strains, was excluded from the analysis.

To assess the clinical relevance of resistance-conferring hits from the screen (shown as red dots in Fig. 2B), we analyzed 49,481 publicly available clinical Mtb genomes^53^ to determine whether these genes exhibit signatures of selection. Strikingly, the uncharacterized gene *rv2571c* showed strong evidence of diversifying selection across its entire coding sequence, ranking second only to *pncA* among resistance-associated genes identified in the screen (**Fig. 2C,D, Data S2**). Phylogenetic analysis of a representative subset of 5,413 clinical Mtb isolates (**Table S2**) revealed repeated, convergent emergence of nonsynonymous and indel mutations in *rv2571c* across multiple Mtb lineages (**Fig. 2E, Data S2**), consistent with adaptive selection in clinical Mtb isolates. *rv2571c* variants were significantly enriched in drug-resistant (DR) clinical Mtb isolates (p=1.18×10⁻⁷), with particularly strong associations with rifampicin resistance (RIF-R, p=1.15×10⁻⁸) and isoniazid resistance (INH-R, p=0.00114) (**Fig. 2F, Supplemental Data 2**). This association pattern reflects the multiple resistance mutations present in many multidrug-resistant (MDR) Mtb isolates^55,56^. Interestingly, loss-of-function mutations in *pncA* were significantly less frequent in isolates harboring *rv2571c* coding variants (4.9%) than in *rv2571c* wild-type strains (15.2%) (**Fig. 2G**), indicating that mutations in these two genes do not typically arise sequentially. Instead, *rv2571c* appears to represent a potentially *pncA*-independent genetic route to PZA resistance.

Guided by these genomic and functional data, we generated a deletion mutant (Δ*rv2571c*) and confirmed that loss of *rv2571c* confers increased resistance to PZA *in vitro* (**Fig. 2H**). This resistance was specific to PZA, as Δ*rv2571c* retained wild-type susceptibility to other first-line antitubercular drugs (**Fig. S4,5B**). Complementation with *rv2571c* expressed from its native promoter fully restored PZA susceptibility, whereas overexpression (∼5-fold) of *rv2571c* from the *hsp60* promoter resulted in hypersusceptibility to both PZA and its active metabolite POA (**Fig. 2H**; **Fig. S5A,C**). We observed similar patterns of susceptibility to structural and functional analogues of PZA and POA, including nicotinamide, nicotinic acid, and benzoic acid (**Fig. S5D**). As previously reported, these compounds inhibited Mtb growth only under acidic conditions (**Fig. S5E**), confirming their pH-dependent activity and implicating *rv2571c* broadly in this class of antimycobacterial compounds^6,8,57,58^. Finally, we assessed the physiological relevance of *rv2571c*-mediated resistance in infection models. The Δ*rv2571c* mutant exhibited increased resistance to PZA during macrophage infection *ex vivo* (**Fig. 2I**) and in mice (**Fig. 2J**; **Fig. S5F**), demonstrating that *rv2571c* contributes to PZA susceptibility *in vivo*.

Together, these results identify *rv2571c* as a previously unrecognized, clinically relevant determinant of PZA susceptibility, distinct from canonical *pncA*-mediated mechanisms, and establish its functional importance across *in vitro*, *ex vivo*, and *in vivo* contexts.

### Defining the mutational landscape and structural basis of Rv2571c-mediated PZA susceptibility

Secondary structure analysis classifies Rv2571c as a multi-pass transmembrane protein whose transmembrane region corresponds to a Fusaric acid resistance protein family (FUSC) domain found in predicted integral membrane transport proteins. Although sequence homologs of Rv2571c are restricted to bacteria, structure-based homology searches revealed unexpected and striking similarity to plant aluminum-activated malate transporters (ALMTs). In particular, *Arabidopsis thaliana* ALMT1 and related ALMT family members emerged as top structural matches, despite minimal primary sequence conservation^59–62^. ALMTs are homodimeric ion channels that conduct organic and inorganic anions through a central pore^61,63^. This strong structural correspondence led us to hypothesize that Rv2571c similarly assembles as a homodimeric channel. AlphaFold-Multimer predictions^64,65^ support this model and reveal an architecture closely resembling ALMT channels, with six transmembrane helices (TM1–TM6) forming a transmembrane domain (TMD) and a C-terminal cytosolic domain (CTD) composed of six α-helices (H1–H6) (**Fig. 3A**; **Fig. S6A**). Dimerization of the TMDs is predicted to create a central pore analogous to that observed in ALMT structures, suggesting a conduit for small anionic metabolites (**Fig. 3A**; **Fig. S6B**).

**Fig. 3.**
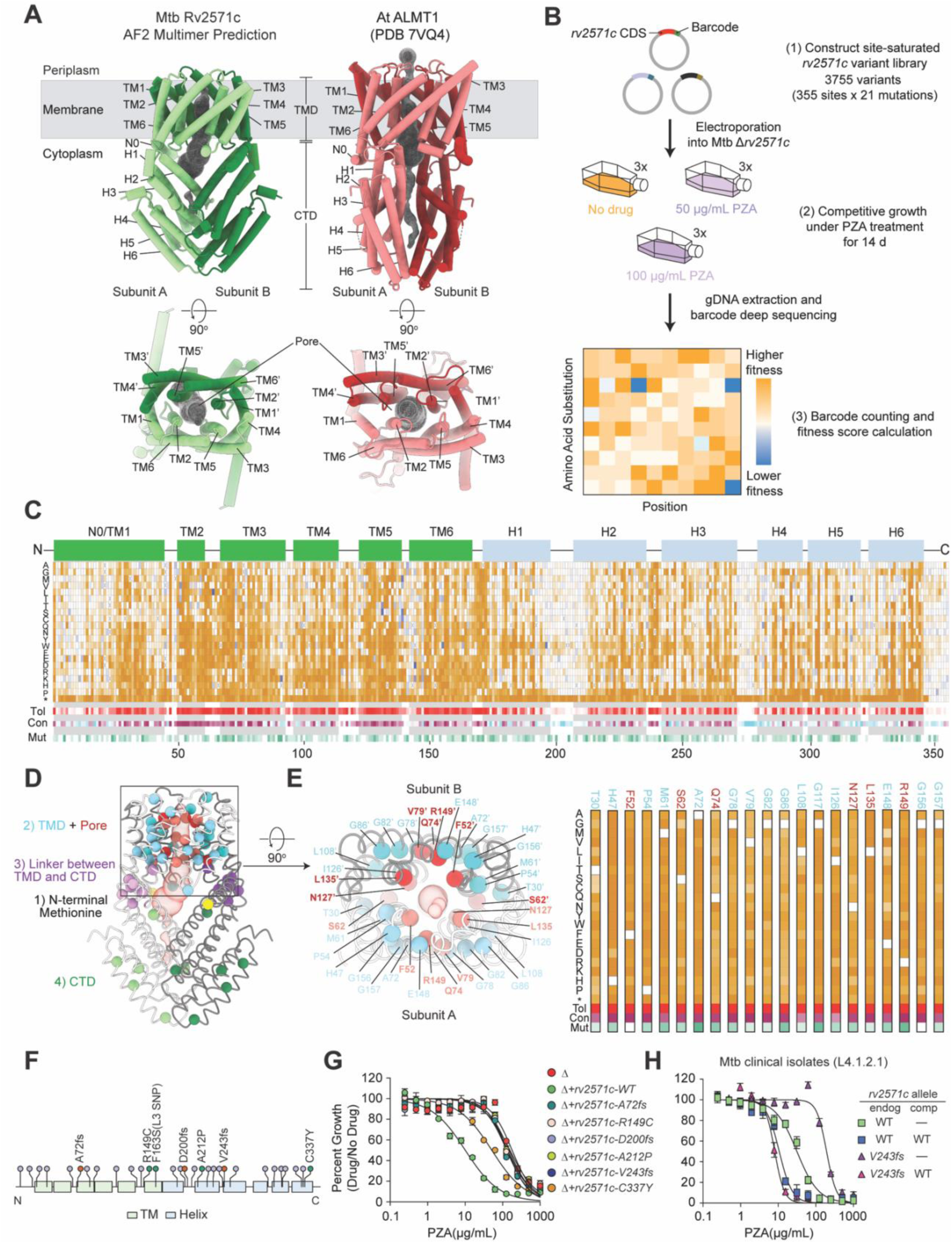
Structural and mutational analysis of Rv2571c-mediated PZA susceptibility. (A) Predicted structure of an Mtb Rv2571c homodimer generated by AlphaFold2 Multimer (left) compared with the cryo-EM structure of *Arabidopsis thaliana* (At) ALMT1 (PDB: 7VQ4) (right). Structures are rendered as cylinders/stubs. Rv2571c protomers are colored pale green and forest green; ALMT1 protomers are colored pale red and dark red. Top row: side view of the dimers; bottom row: top-down view. The grey box indicates the approximate membrane boundaries and cytosolic region. Domains (TMD, transmembrane domain; CTD, C-terminal domain), transmembrane helices (TM), and cytosolic helices (H) are labeled. Putative pores of Rv2571c homodimer and ALMT1 calculated from MOLE are rendered as a grey mesh. (B) Workflow for deep mutational scanning (DMS) of *rv2571c*. A barcoded library containing substitutions at 355 sites within *rv2571c* (21 possible substitutions per site) was cloned into a mycobacterial integrating plasmid. The library was electroporated into Mtb Δ*rv2571c* and subjected to selection in the absence or presence of PZA at two different concentrations (no drug, 50 µg/mL, 100 µg/mL). After selection, genomic DNA was extracted, barcodes were PCR-amplified and deep-sequenced, and fitness scores were calculated for each variant. (C) Heatmaps showing functional effects (z-scores) of nearly all possible amino acid substitutions in Rv2571c under 50 µg/mL PZA selection. The secondary-structure topology of Rv2571c is shown at the top, with green boxes indicating transmembrane (TM) and light blue boxes for cytosolic (H) helices. The main functional-score heatmap displays z-scores for each substitution (rows) across all positions (columns). Control “stop” codon is represented by “*”; gold indicates loss-of-function (PZA resistance) and blue indicates gain-of-function (PZA sensitivity) variants. Also shown are heatmaps for mutational intolerance scores (Tol, tolerant (white) to intolerant (red)), conservation scores (Con, cyan (variable) to white to purple (conserved)), and frequency of mutations observed in clinical isolates (Mut, white (low) to green (high)). (D) Predicted structure of the Rv2571c dimer rendered as licorice/ovals cartoons. The putative pore is rendered as a red surface (50% transparency). Alpha-carbon atoms of the most mutationally intolerant residues (n=36) are shown as spheres and color-coded by structural cluster: blue/cyan (transmembrane domain), red/salmon (pore region), pink/purple (linker region), yellow (N-terminal methionine), pale green/forest green (C-terminal domain). The transmembrane domain is highlighted with a black box. (E) Mutationally intolerant residues in the transmembrane domain cluster within the putative pore of the Rv2571c dimer. Left panel: zoomed-in view of the transmembrane domain boxed in panel D. Structure is rendered as licorice/ovals cartoons: subunit A is white, subunit B is grey, and the putative pore is rendered as a red surface (50% transparency). Mutationally intolerant residues are shown as spheres: blue/cyan represents residues making up the transmembrane domain, and red indicates residues that line the putative pore. Right panel: Heatmaps of functional scores, mutational tolerance, conservation, and frequency of clinical mutations for the displayed residues. (F) Diagram of the Mtb *rv2571c* open reading frame showing the 30 most common coding variants observed in our clinical strain genome database. Specific mutations tested in panel G are annotated. (G) PZA dose-response curves of Mtb Δ*rv2571c* complemented with *rv2571c* WT or clinically observed *rv2571c* variants. (H) PZA dose-response curves of two clinical strains (lineage L4.1.2.1) harboring either a WT (green square) or an *rv2571c* LOF variant (*rv2571-V243fs*, purple triangle). A WT copy of *rv2571c* was introduced in trans (*+rv2571c-WT*) in each clinical strain (navy square, and burgundy triangle, respectively). G and H are means ± s.d of an experiment performed in triplicate and representative of three independent experiments.

To functionally interrogate Rv2571c and identify residues critical for PZA susceptibility, we performed deep mutational scanning (DMS) across the entire protein^66,67^. We constructed a barcoded, site-saturated library encoding all possible single–amino acid substitutions, including stop and synonymous variants, across the 355-residue *rv2571c* coding sequence, covering ∼95% of all possible variants. The variant library was integrated into Mtb Δ*rv2571c* and subjected to competitive growth in acidic HBO medium, with or without PZA (50 or 100 µg/mL; **Fig. 3B**). Variant fitness was quantified by barcode sequencing and analyzed using a negative binomial regression framework to derive fitness z-scores (**Data S3**). Biological replicates showed high reproducibility, and the 50 and 100 µg/mL PZA selections yielded highly similar fitness landscapes (**Fig. S7A,B**). Because the 50 µg/mL dataset provided optimal barcode coverage, clear separation of premature stop variants, and matched the conditions of the CRISPRi screen, we focused subsequent analyses on this condition (**Fig. S7C,D**).

As expected, disruption of Rv2571c function conferred a strong fitness advantage under PZA selection: mutation of the start codon or introduction of premature stop codons at most positions resulted in increased fitness in the presence of PZA (**Fig. 3C**). To systematically define residues essential for Rv2571c function, we integrated DMS-derived fitness scores with conservation metrics and mutation frequencies from our clinical Mtb genome database. This analysis revealed strong concordance between mutational intolerance, evolutionary conservation, and enrichment of variants in clinical isolates (**Fig. 3C**; **Fig. S7E,F**). In total, we identified 36 residues that were completely intolerant of mutation under PZA selection (**Fig. 3D**). These residues cluster within four key regions: (i) the N-terminal methionine; (ii) the TMD; (iii) the linker connecting the TMD and CTD; and (iv) structured helices of the CTD. In contrast, residues tolerant of mutation were largely confined to loop regions, consistent with structural flexibility in these regions.

Mapping intolerant residues onto the AlphaFold-Multimer structure revealed that the majority are in the TMD (**Fig. 3D**), contributing to the overall fold of this domain and lining the central pore formed by the two Rv2571c subunits (**Fig. 3E**). Notably, ALMT channels contain two conserved arginine residues within TM3 and TM6 that coordinate malate within the pore^59,61,62^. While Rv2571c lacks the TM3 arginine, it retains the homologous TM6 residue R149. R149 is completely intolerant of mutation, highly conserved, positioned directly within the predicted pore, and harbors variants in DR clinical Mtb isolates (**Fig. 3E**), implicating it in substrate coordination or conductance. Collectively, these structural and mutational data support the conclusion that Rv2571c functions as a homodimeric channel protein.

To experimentally validate the functional consequences of clinically observed *rv2571c* variants, we expressed selected frameshift and missense alleles in Δ*rv2571c* Mtb (**Data S2**; **Table S3**) and assessed PZA susceptibility. All predicted loss-of-function variants, including the pore residue mutant R149C, failed to restore PZA susceptibility, consistent with DMS predictions (**Fig. 3F,G**). In contrast, the lineage 3–specific SNP F163S, which is fixed across lineage 3 strains and not associated with PZA resistance, retained function, in agreement with the DMS data (**Fig. S5G**).

Finally, we examined a clinical Mtb isolate harboring a V243 frameshift mutation in *rv2571c*. This isolate exhibited greater PZA resistance than a phylogenetically matched strain carrying wild-type *rv2571c* (**Fig. 3H**). Complementation with wild-type *rv2571c* restored PZA susceptibility, confirming that loss of *rv2571c* function was the determinant of PZA resistance in this clinical isolate.

Together, these data define the mutational landscape of Rv2571c, establish its structural similarity to ALMT anion channels, and demonstrate that loss of Rv2571c function, through diverse and clinically observed mutations, constitutes a novel, mechanistically distinct route to PZA resistance in Mtb.

### Rv2571c enables α-ketoglutarate export

Although Rv2571c is predicted to function as an anion channel based on strong structural homology to the ALMT protein family, its physiological substrate remains unknown. Prior work suggested a potential role in drug transport, based on the resistance of *rv2571c* loss-of-function mutants to arylamide compounds^68^. We therefore tested whether Rv2571c influences PZA or POA transport. Contrary to this hypothesis, targeted metabolomic analyses revealed comparable intracellular and extracellular levels of PZA and POA in wild-type and Δ*rv2571c* Mtb (**Fig. S8A, B**), indicating that PZA resistance in Δ*rv2571c* is not attributable to altered PZA or POA accumulation.

To identify the physiological substrate of Rv2571c, we expressed *rv2571c* in *Mycobacterium smegmatis* (Msmeg), which lacks a native homolog. Unexpectedly, induction of *rv2571c* expression led to the formation of a white precipitate on solid medium (**Fig. S8C**) and was accompanied by localized acidification of the surrounding agar (**Fig. 4A**). Untargeted metabolomic profiling of surrounding media identified 13 features significantly enriched upon *rv2571c* expression (**Fig. 4B**, **Data S4**). Among these, two of the most abundant signals (ranked first and fifth) corresponded to ions at m/z 101.024 and 145.013, annotated as succinic semialdehyde (SSA) and α-ketoglutarate (αKG), respectively. Tandem MS analysis demonstrated that the SSA signal arises from in-source fragmentation of αKG, consistent with previous reports^69,70^, establishing αKG as the dominant exported metabolite (**Fig. 4C**; **Fig. S8D**). Validating this result, heterologous expression of wild-type *rv2571c*, but not the loss-of-function R149C variant, led to robust αKG accumulation in liquid culture and acidification of solid medium (**Fig. S8E,F**).

**Fig. 4.**
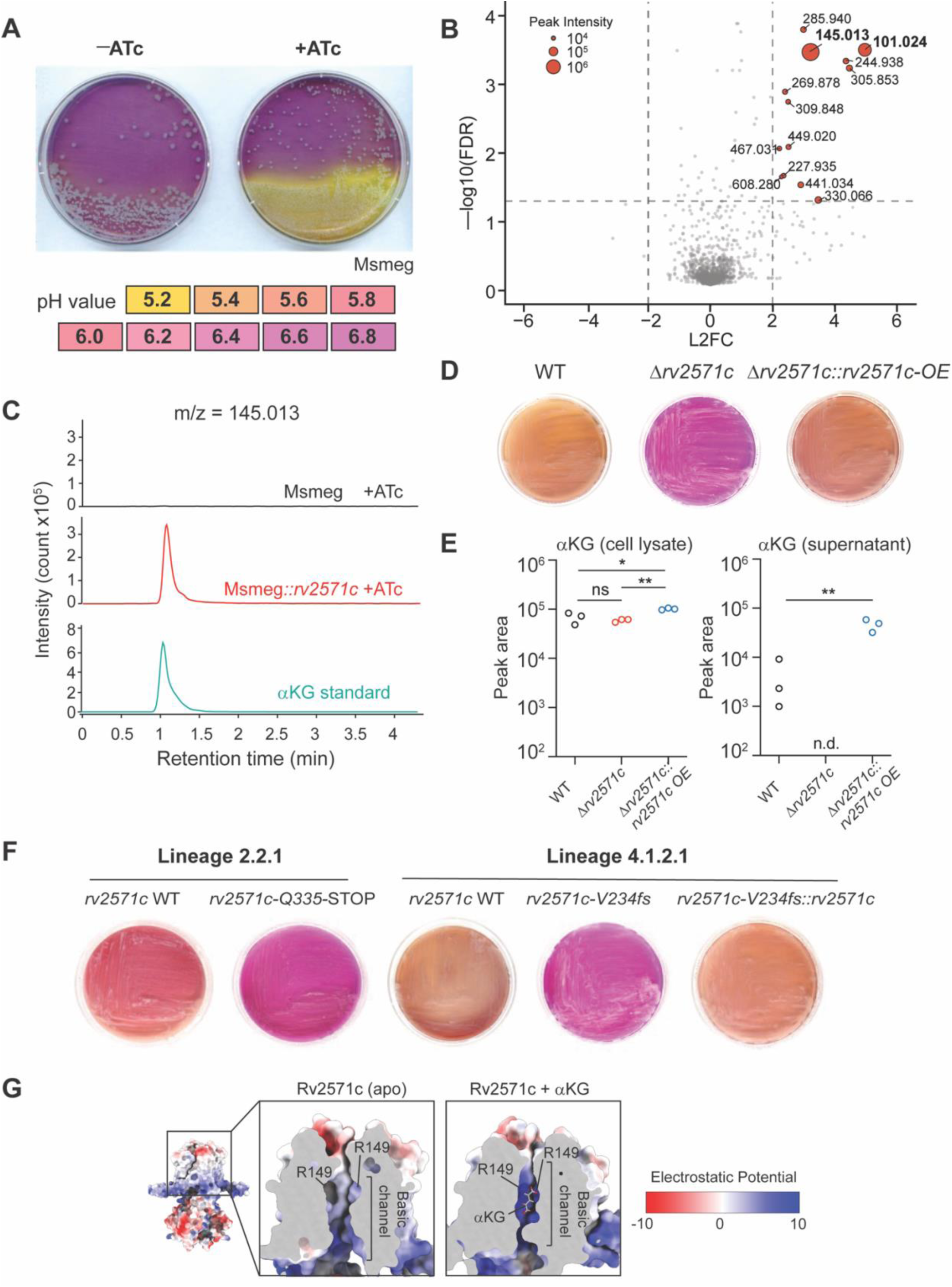
Rv2571c facilitates the export of alpha-ketoglutarate (αKG). (A) Heterologous expression of *rv2571c* in Msmeg alters medium pH. Msmeg transformed with an ATc-inducible *rv2571c* construct was cultured on 7H10 agar containing the pH indicator chlorophenol red (CPR), in the presence or absence of ATc. Representative changes in medium color corresponding to pH values ranging from 5.2 to 6.8 are shown. Pictures are representative of three biological replicates and were captured after 3 days of incubation at 37°C. (B) Volcano plot showing differentially abundant features from untargeted metabolomic analysis of agar media conditioned by WT Msmeg and Msmeg expressing *rv2571c*. Dashed lines indicate significance thresholds (−log10 FDR > 1.33 and |L2FC| > 2). Selected m/z features are annotated. Features at m/z 145.013 and 101.024 (highlighted in red) correspond to αKG and succinic semialdehyde (SSA, see **Fig. S8D**). (C) Extracted ion chromatograms showing signals corresponding to deprotonated αKG (m/z = 145.013) in agar media conditioned as in panel B. The signal detected in Msmeg::*rv2571c*–conditioned media co-elutes with an αKG standard. (D) Indicated Mtb strains were cultured on 7H10 agar containing CPR. Pictures are representative of three biological replicates and were captured after 30 days of incubation at 37°C. (E) αKG pool size in cell lysates (left) and culture supernatants (right) from the indicated Mtb strains. Data represent means ± s.d of one experiment performed in triplicate and are representative of at least two independent experiments. Statistical significance was calculated with Student’s *t*-test; *, *P* < 0.05, **, *P* < 0.01. (F) Sublineage-matched clinical Mtb isolates harboring either WT or LOF *rv2571c* alleles were cultured on 7H10 agar containing CPR. Complementation of the L4 strain demonstrates causality. Pictures are representative of three biological replicates and were captured after 30 days of incubation at 37°C. (G) AF2 Multimer prediction for Rv2571c (left) rendered as a molecular surface and colored by electrostatic potential in ChimeraX. Insets show a zoom-in slice through the transmembrane domain indicated by the black box. Left inset shows no substrate bound, and right inset shows the predicted structure of αKG-bound state from DynamicBind.

We next examined whether Rv2571c similarly enables αKG export in Mtb. Wild-type Mtb acidified its surrounding medium in an Rv2571c-dependent manner (**Fig. 4D**). Metabolomics confirmed that deletion of *rv2571c* abolished αKG secretion into the culture supernatant, while intracellular αKG levels were not affected **(Fig. 4E**). Importantly, clinical Mtb isolates harboring *rv2571c* loss-of-function mutations as well as Δ*rv2571c* strains expressing DMS-validated LOF variants, failed to acidify solid medium (**Fig. 4F**; **Fig. S9A**), establishing a strong correlation between αKG export capacity, *rv2571c* genotype, and PZA resistance. In support of a direct transport role, electrostatic surface analysis of the Rv2571c homodimer revealed a centrally positioned basic pore capable of accommodating the dianionic αKG molecule (**Fig. 4G**).

### Rv2571c-dependent αKG export potentiates PZA-induced cytoplasmic acidification

PZA has been proposed to act through multiple mechanisms, including inhibition of trans-translation (RpsA)^25^, fatty acid synthesis (FAS-I)^26^, and coenzyme A biosynthesis (PanD)^27,29^. However, our chemical–genetic screen did not support these target-based models. Instead, our data are consistent with a model in which PZA activity depends on an acidic extracellular milieu and results from drug-induced cytoplasmic acidification^8,30^. In this framework, PZA is converted by PncA into POA, which then shuttles protons from the acidic extracellular space into the near-neutral bacterial cytoplasm, ultimately driving lethal cytoplasmic acidification.

Given this model, we examined whether Rv2571c-dependent αKG export influences PZA susceptibility by modulating cytoplasmic pH. Consistent with this hypothesis, we observed that overexpression of *rv2571c* increases extracellular αKG accumulation and causes a pronounced growth defect under acidic conditions (pH < 5.5; **Fig. S9B-D**). These observations suggest that excessive αKG export is deleterious in acidic environments and raise the possibility that Rv2571c-dependent αKG export influences cytoplasmic pH homeostasis and PZA susceptibility.

To test this hypothesis, we measured intracellular pH using a validated pH-GFP reporter^8,39,71^. As expected, PZA caused dose-dependent cytoplasmic acidification, whereas growth-inhibitory concentrations of isoniazid did not alter intracellular pH, confirming that cytoplasmic acidification is specific to PZA activity (**Fig. S10A,B**). Strikingly, Δ*rv2571c* was significantly more resistant to PZA- and POA-induced cytoplasmic acidification than WT (**Fig. 5A-D, Fig. S10E**). Conversely, the *rv2571c* overexpression strain, which exports elevated levels of αKG and grows poorly at low pH, exhibited a lower basal intracellular pH even in the absence of PZA (**Fig. 5A**). Consistent with the broader resistance profile of Δ*rv2571c* to nicotinamide, nicotinic acid, and benzoic acid (**Fig. S5D**), these compounds also induced cytoplasmic acidification in WT Mtb, an effect that was attenuated in Δ*rv2571c* (**Fig. S10E-H**). Together, these results indicate that Rv2571c-dependent αKG export sensitizes Mtb to cytoplasmic acidification across this class of pH-dependent antimycobacterial compounds.

**Fig. 5.**
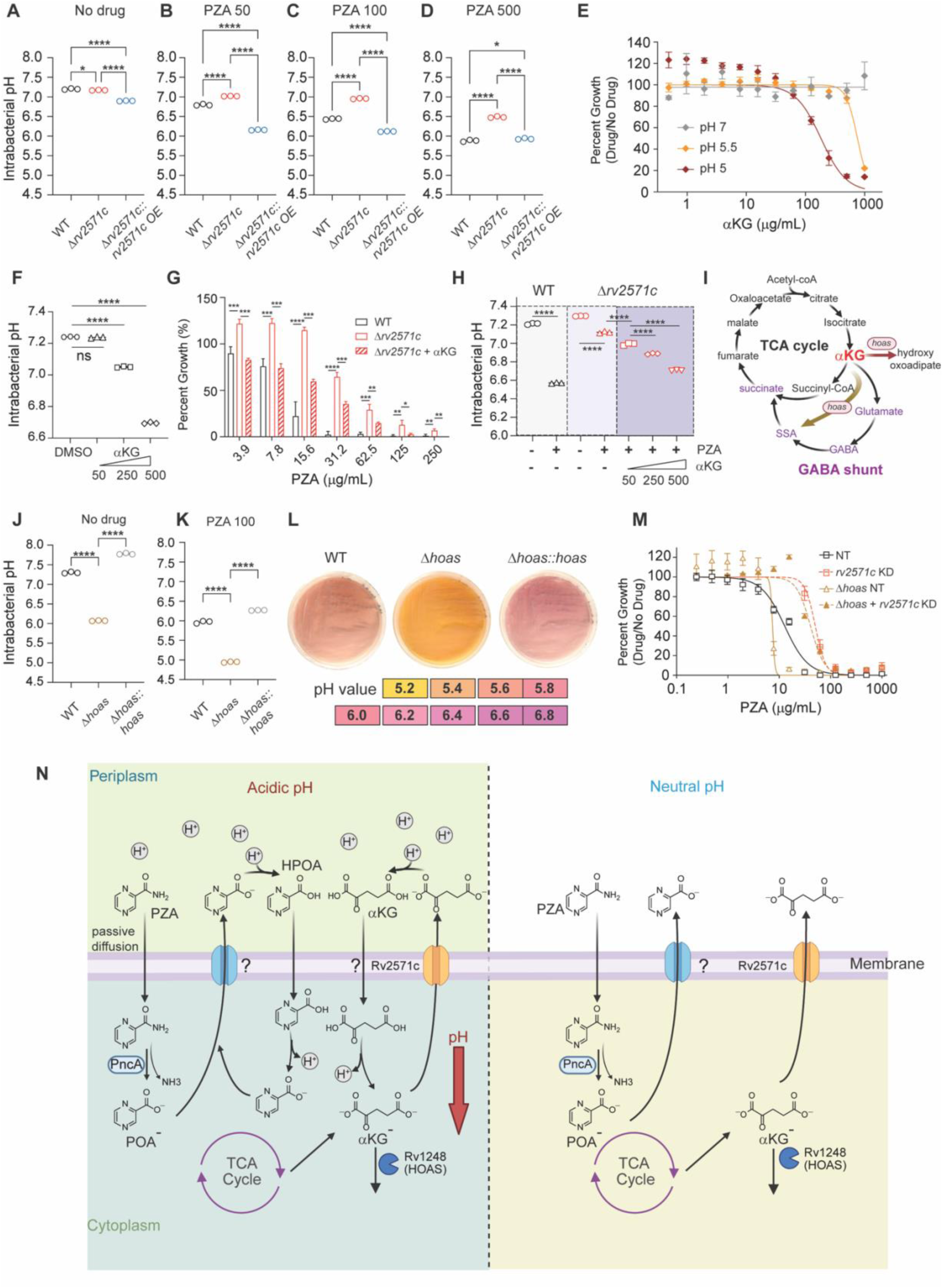
Rv2571c-dependent αKG export modulates PZA-mediated cytoplasm acidification. (A–D) Intrabacterial pH measurement of the indicated Mtb strains grown without drug (A) or in the presence of 50 µg/mL (B), 100 µg/mL (C), or 500 µg/mL (D) PZA. (E) Growth inhibition of Mtb in the presence of increasing concentrations of αKG in HBO media at pH 7.0, 5.5, or 5.0. (F) Intrabacterial pH measurement of WT Mtb in the presence of DMSO or increasing concentrations of exogenous αKG (50, 250, or 500 µg/mL). (G) Growth of indicated Mtb strains in the presence of αKG supplementation (250 μg/mL) across a range of PZA concentrations. (H) Intrabacterial pH measurement of WT and Δ*rv2571c* Mtb treated (+) or untreated (–) with PZA (100 µg/mL). Δ*rv2571c* strains treated with PZA were additionally supplemented with increasing concentrations of αKG (250, 500, and 750 μg/mL). (I) Schematic of major αKG metabolic pathways in Mtb, highlighting the role of HOAS at the interface of the TCA cycle and the GABA shunt. (J–K) Intrabacterial pH measurement of the indicated Mtb strains grown in HBO medium at pH 5 in the absence or presence of 100 µg/mL PZA. (L) Representative images of WT, Δ*hoas*, and complemented Δ*hoas*::*hoas* Mtb plated on CPR-containing 7H10 plates. Reference color scale indicating varying pH values is shown. Pictures are representative of three biological replicates and were captured after 30 days of incubation at 37°C. (M) Dose-response curves for WT, Δ*hoas*, and corresponding CRISPRi knock-down (KD) strains treated with PZA. NT, non-targeting sgRNA control. (N) Proposed model linking Rv2571c-mediated αKG export to PZA susceptibility via cytoplasm acidification. PZA enters the cytoplasm by passive diffusion and is hydrolyzed by PncA to generate pyrazinoic acid (POA⁻). Under acidic extracellular conditions (left), exported POA⁻ becomes protonated to form HPOA, which passively diffuses back into the cell, where it dissociates and releases a proton, driving cytoplasmic acidification through a POA cycling mechanism. Rv2571c facilitates the export of αKG, which, under acidic extracellular conditions, becomes protonated in a manner analogous to pyrazinoic acid (POA). Protonated αKG can re-enter the bacterium by passive diffusion, thereby potentiating POA-mediated cytoplasmic acidification. In parallel, HOAS (Rv1248c) consumes αKG and modulates intracellular αKG availability. At neutral extracellular pH (right), the reduced proton concentration limits the formation of protonated POA and αKG, thereby restricting their re-entry and attenuating intrabacterial acidification. This model was created using BioRender.com. In panels A-H, J-K and M, data represent means ± s.d of one experiment performed in triplicate and representative of at least two independent experiments; statistical significance was determined by one-way ANOVA followed by Tukey’s post-hoc test. *, adj-*P* < 0.05, **, adj-*P* < 0.005, ***, adj-*P* < 0.0005, ****, adj-*P* < 0.0001.

αKG export could sensitize Mtb to cytoplasmic acidification through multiple mechanisms, including indirect effects on membrane electrochemical properties or through the direct activity of extracellular αKG itself. Aligned with the latter possibility, exogenous αKG inhibited Mtb growth in a pH-dependent manner, with strong inhibition observed at pH ≤ 5.5 but minimal effects at neutral pH (**Fig. 5E**). αKG supplementation also caused cytoplasmic acidification, although less efficiently than PZA (**Fig. 5F**). Importantly, the addition of exogenous αKG restored both PZA susceptibility and cytoplasmic acidification to Δ*rv2571c* Mtb, indicating that loss of αKG export underlies PZA resistance in this mutant (**Fig. 5G,H**). αKG inhibited growth and induced cytoplasmic acidification without reducing the pH of the surrounding medium, indicating that its effects are not mediated by bulk extracellular acidification.

To further link αKG metabolism to PZA susceptibility, we interrogated our CRISPRi screen data (**Data S1**) for genes whose knockdown sensitized Mtb to PZA and identified *hoas* (*rv1248c*), encoding a 2-hydroxy-3-oxo-adipate synthase, a key αKG-consuming enzyme^72,73^ (**Fig. 5I)**. Consistent with prior reports^73^, Δ*hoas* accumulated αKG intracellularly and extracellularly, acidified solid media more strongly than WT, phenocopied *rv2571c* overexpression, displaying impaired growth at acidic pH, and reduced intracellular pH even in the absence of drug (**Fig. 5J-L**; **Fig. S11A,B**). As predicted by the CRISPRi screen, Δ*hoas* was hypersensitive to PZA-induced cytoplasmic acidification and killing (**Fig. 5K**; **Fig. S11C**). Crucially, CRISPRi knockdown of *rv2571c* in the Δ*hoas* background reduced αKG export (**Fig. S 11D**) and reversed PZA hypersusceptibility (**Fig. 5M**), establishing that Rv2571c-dependent αKG export is required for the heightened PZA sensitivity caused by loss of HOAS.

Finally, because PZA is active against non-replicating Mtb, we examined whether αKG-mediated modulation of PZA susceptibility persisted under non-replicating conditions. Both Δ*rv2571c* and Δ*hoas* retained their respective PZA resistance and hypersensitivity phenotypes in non-replicating bacteria, demonstrating that αKG export influences PZA activity independently of bacterial replication or carbon availability (**Fig. S12**).

Collectively, these data support a model in which Rv2571c-dependent export of αKG potentiates PZA-induced cytoplasmic acidification specifically under acidic conditions. These findings uncover a previously unrecognized metabolic dimension of PZA’s mechanism of action and identify Rv2571c as a critical determinant of pH-dependent drug susceptibility. Based on its function, we propose renaming Rv2571c as AKGC (α-ketoglutarate channel).

## DISCUSSION

PZA is a cornerstone of modern TB therapy, yet its mechanism of action and clinically relevant resistance mechanisms beyond disrupting PncA activity have remained difficult to define, owing in part to the experimental challenges inherent to studying PZA. Here, by developing a lipid-rich, acidic culture system that robustly captures PZA’s bactericidal activity, we establish three core conclusions. First, acidic pH is indispensable for PZA-mediated killing. Second, integrating this system with CRISPRi chemical-genetics and comparative genomics identifies LOF mutations in *rv2571c* as a major, clinically relevant determinant of PZA susceptibility. Third, mechanistic dissection supports a model in which Rv2571c functions as an anion channel that enables export of αKG, and that αKG export sensitizes Mtb to PZA by potentiating PZA/POA-driven cytoplasmic acidification. Together, these findings connect ion-channel–mediated metabolite flux, central carbon metabolism, and cytoplasmic pH homeostasis to PZA susceptibility, with implications for improving resistance detection and guiding the development of treatment-shortening PZA-like therapeutics.

LOF mutations in *rv2571c* are likely the “missing mechanism” for phenotypic PZA resistance in many clinical Mtb isolates that lack *pncA* mutations^17,20^. *rv2571c* mutations appear to confer a largely *pncA*-independent route to PZA resistance; less frequently, they may also facilitate subsequent acquisition of higher-level resistance through *pncA* inactivation. Although *rv2571c* loss confers a lower level of resistance than canonical *pncA* inactivation, this effect is nonetheless clinically meaningful and has substantial impact during infection (**Fig. 2 I-J**). *rv2571c* LOF alleles are associated with MDR-TB outbreaks across multiple geographic regions^74–78^ and continue to arise during the course of TB therapy^79^, indicating ongoing positive selection in patients, as also reported in several independent studies^76,80,81^. While such resistance might appear surmountable by increasing PZA dose, this strategy is unlikely to be viable in practice: PZA dosing is limited by tolerability^82^ and inter-individual variability in PZA pharmacokinetics may leave a significant fraction of patients below exposures required for therapeutic benefit^83,84^.

Mechanistically, our data support a model–first proposed by Zhang and colleagues–in which PZA functions less like a classical, target-specific antibiotic and more like a weak-acid “pro-preservative,” whose bactericidal activity is revealed only in acidic extrabacterial environments that permit repeated cycles of POA protonation and re-entry (**Fig. 5N**). In this framework, PZA is converted to POA⁻ in the cytoplasm by PncA and, via an as-yet undefined pathway, is exported from the cell; under acidic conditions, POA⁻ becomes protonated (HPOA), re-enters the bacterium, and dissociates to release protons, thereby driving cytoplasmic acidification through a cycling mechanism.

The central advance described here is the discovery that αKG export facilitated by Rv2571c (AKGC) functionally links central carbon metabolism to this weak-acid cycling mechanism. Although several mechanistic possibilities could explain this connection, our favored model is that exported αKG becomes protonated in acidic environments in a manner analogous to POA, and that protonated αKG can then re-enter the cell to further amplify cytoplasmic acidification. This framework parsimoniously explains why loss of Rv2571c diminishes PZA-induced cytoplasmic acidification, why exogenous αKG restores both acidification and drug susceptibility in Δ*rv2571c* strains, and why perturbing αKG consumption (for example, in Δ*hoas*) shifts basal intracellular pH and creates PZA hypersusceptibility in a manner dependent on Rv2571c.

These findings help reframe longstanding debates surrounding PZA’s mechanism of action. Numerous target-based models have been proposed^11–13^, including those implicating CoA biosynthesis pathways (most notably via PanD)^27–29^, and many of these observations are compelling within specific experimental contexts. In contrast, earlier models invoking inhibition of fatty acid synthesis or trans-translation have not been supported by subsequent experiments^85,86^. Our results do not contradict prior studies, but they do argue that in a lipid-rich, acidic culture system that more closely mimics infection conditions, cytoplasmic acidification is tightly linked to growth inhibition and emerges as the dominant physiological axis of PZA activity. The observation that loss of *rv2571c* confers protection not only against PZA but also against chemically related, pH-dependent aromatic acids (such as nicotinate and benzoate) is more consistent with a shared biophysical mechanism of acidification than with shared inhibition of a common protein target. Moreover, the repeated selection for *rv2571c* LOF in TB patients receiving first-line therapy^74–78,80,81^—contrasting with the absence of comparable selection signatures for several other proposed targets like *panD*—supports the conclusion that αKG-linked physiology and cytoplasmic acidification represent a clinically relevant route to PZA failure in humans.

Important caveats remain. Although cytoplasmic acidification and growth inhibition are tightly correlated in our system, we cannot formally distinguish whether acidification is the direct cause of bacterial death or instead a closely linked consequence of perturbing another essential process. Weak acids are known to exert multiple effects on bacteria, including cytoplasmic acidification, collapse of membrane potential, inhibition of substrate transport, and broader membrane-associated stress.

There are also important mechanistic uncertainties regarding the channel itself. Although our data strongly support αKG export mediated by Rv2571c/AKGC, definitive identification of substrates will ultimately require biochemical reconstitution and direct flux measurements. Moreover, αKG conductance does not preclude the possibility that AKGC transports other metabolites. The observation that plant ALMT homologues conduct the related TCA cycle intermediate malate is nevertheless consistent with αKG as a physiologically relevant substrate. Finally, because ALMT-family channels are known to exhibit complex regulation by ligands, lipids, and voltage, elucidating the factors that gate AKGC opening and closing—and how this regulation is integrated with infection-state metabolism—will be essential for developing a complete physiological model.

Expression of *rv2571c*, particularly overexpression in Msmeg, results in extracellular acidification on solid media. The source of the protons accumulating in the medium in response to Rv2571c expression remains unclear. Notably, acidification is observed on solid medium but not in liquid culture, possibly reflecting the greater bacterial biomass achieved on solid media. One potential source of protons is the export of the monoprotonated form of αKG. Although this species constitutes only ∼0.34% of intracellular αKG at pH 7^87^, it could accumulate at high cell density and release protons upon dissociation in the extracellular environment. This scenario would presumably require AKGC to transport both the unprotonated and monoprotonated forms of αKG. Alternatively, proton export may occur secondarily to maintain membrane electrochemical balance during αKG efflux. Such coupling has precedent: in plants, secretion of organic acids—including via ALMT family channels—is frequently accompanied by proton export through H⁺-ATPases to preserve the membrane electrochemical gradient ^88–92^. In mycobacteria, this compensation would likely occur through increased respiratory proton pumping rather than reversal of the ATP synthase, as the ATPase is not thought to operate efficiently in reverse^93^. Thus, enhanced electron transport chain activity could serve to restore membrane electrochemical balance during sustained αKG efflux, indirectly linking anion export to extracellular acidification.

Our findings also raise a basic physiological question: why would Mtb export αKG? αKG, along with other TCA cycle intermediates such as malate, isocitrate, and succinate, has been reported to be secreted by Mtb under standard growth conditions and in response to diverse stresses, including hypoxia, iron limitation, and antibiotic exposure. One possibility is that αKG export reflects metabolic “overflow” or redox balancing, with electrogenic anion export coupled to maintenance of intracellular homeostasis. The fact that *rv2571c* LOF confers PZA resistance in macrophages (**Fig. 2I**), mice^94^ (**Fig. 2J, Fig. S5F**), and presumably in humans^74–78,80,81^ (**Fig. 2C-J**), implies that αKG export likely occurs in the host environment, potentially into the phagosomal lumen. Parsing when and where AKGC is active *in vivo* should help connect this channel to infection-state metabolism and to the selective conditionality of PZA.

From a translational perspective, our findings suggest both opportunities and cautions for improving TB therapy. One strategy is the development of next-generation PZA-like weak-acid prodrugs with physicochemical properties optimized to support efficient proton cycling in host-relevant environments. The use of HBO medium together with cytoplasmic acidification reporters may facilitate the identification and optimization of such compounds. Because systemic administration of POA is ineffective^95^ and toxicity must be avoided, such compounds, like PZA, may need to be bioactivated and selectively concentrated within Mtb. That said, the weak acid aspirin—rapidly converted to salicylic acid by host esterases in humans—induces cytoplasmic acidification in Mtb akin to POA^96,97^, which may contribute to the reported benefit of aspirin as an adjunctive agent in TB patients^98,99^. A second, complementary approach is rational potentiation: our genetic data highlight nodes in central carbon metabolism, including αKG-consuming enzymes such as HOAS, as well as other CCM enzymes like PEPCK (coded by *pckA*), whose inhibition may synergize with PZA-like chemistry, potentially providing a “two-for-one” advantage given their established roles promoting Mtb fitness in the host^73,100^. In parallel, the identification of *blaR* as a sensitizing factor suggests that co-inhibition of respiration or electron transport chain function^101,102^ could further amplify PZA activity, consistent with a close coupling between cellular energetics and acid stress–mediated lethality. Our findings also raise the possibility of directly modulating AKGC activity itself: small-molecule agonists that promote channel opening or increase conductance could amplify αKG export and thereby enhance PZA-induced cytoplasmic acidification. Such a strategy would conceptually parallel chemical agonists like menthol and capsaicin that lower the activation threshold for cold and heat-sensing TRP ion channels, respectively^103,104^. At the same time, an important caution emerges: *rv2571c* appears dispensable for bacterial fitness in humans, implying that therapeutic strategies that rely on AKGC activity for potentiation may exhibit reduced efficacy in the presence of pre-existing or acquired channel loss-of-function mutations.

Finally, our study has immediate implications for diagnostics. PZA drug-susceptibility testing is notoriously difficult, and uncertainty about PZA activity in MDR-TB regimens remains a persistent clinical dilemma^21–24^. By enabling robust PZA activity *in vitro*, the lipid-rich, acidic medium described here offers a practical route to improve the reliability of phenotypic testing. In parallel, incorporation of *rv2571c* into molecular diagnostic frameworks, alongside *pncA* and informed by our DMS data could enhance detection of clinically relevant resistance mechanisms.

In sum, this work resolves key barriers that have long obscured PZA’s mechanism of action. Our findings demonstrate that acidic pH is essential for PZA-mediated killing, identify LOF mutations in *rv2571c* as a major and clinically relevant determinant of PZA resistance, and support a model in which Rv2571c functions as an ion channel whose activity sensitizes Mtb to PZA by amplifying POA-driven cytoplasmic acidification. Together, these findings have direct implications for improving resistance detection and for guiding the rational development of next-generation, treatment-shortening PZA-like therapies.

## ACKNOWLEDGEMENTS

We thank Guanghui Yang, Jiashuai Fan, Bill Schneider, Leon Chan, Nathan Lubock, Nathan Hicks, and Veronique Dartois for helpful discussions and/or comments on the manuscript, and Anthony Castro, Rodrigo Aguilera Olvera and Jialing Wang for technical assistance. We thank Martin Bush and Thomas Walz for their valuable contributions to the early stages of this project and for providing preliminary experimental data. We thank Ruslana Bryk and Carl Nathan for providing the Δ*hoas* Mtb mutant and related strains. During the preparation of this manuscript the authors used Microsoft 365 Copilot (for business/enterprise) in order to improve clarity of language. After using this service, the authors reviewed and edited the content as needed and take full responsibility for the content of the published article.

## AUTHOR CONTRIBUTIONS

Conceptualization: AG, SL, SE, JMR;

Methodology and Investigation: AG, SL, JC, AN, AS, ZAA, KT, VMG, NCP, MAD,KR;

Writing – original draft: AG, SL, SE, JMR;

Writing – review & editing: JC, AN, ZAA, NCP and DS;

Funding acquisition: AG, SE, JMR;

Supervision: SE, JMR

## DATA, CODE, AND MATERIAL AVAILABILITY

All data are available in the manuscript or the supplementary materials

## SUPPLEMENTAL MATERIAL

Materials and Methods

Figs. S1 to S12

Tables S1 to S5

Data S1 to S4

