## Supplemental Figures and Material for "Metabolic control of drug resistance by a mycobacterial ion channel"

**The PDF file includes:**

Materials and Methods

Figs. S1 to S12

Tables S1 to S5

**Other Supplementary Materials for this manuscript include the following:**

Data S1 to S4

**Material and Methods**

**Bacterial strains and reagents**

*Mycobacterium tuberculosis* (Mtb) strains are derivatives of H37Rv unless otherwise noted. The mutant strains ∆*rv2571c*, ∆*hupB* and ∆*blaR* were constructed in Mtb for this study using a recombineering method. The mutant ∆*pckA* and complemented strain (∆*pckA::pckA*) were constructed by us as previously described^40^. The mutant ∆*pncA* as well as its related WT were gifted by Helena I. M. Boshoff (NIH, Bethesda, USA) as previously described^105^. The mutant ∆*hoas* as well as its complemention (*hoas::hoas*) were gifted by Carl Nathan (Weill Cornell Medicine, New York, USA) and previously described^73^. The set of strains WT, ∆*rv2571c*, ∆*rv2571c::rv2571c*-OE and WT, ∆*hoas*, ∆*hoas::hoas* were electroporated with an episomal plasmid harboring a kanamycin resistant cassette and allowing the expression of the pH-GFP reporter downstream the promoter Ptb38 as similarly described^71,96^. The collection of ∆*rv2571c* strains expressing the *rv2571c* variants was constructed for this study (See details in *Generation of Mtb mutants and genetic complementation* for details regarding strain construction). A reference set of Mtb clinical strains was obtained from the Belgian Coordinated Collections of Microorganisms (BCCM)^106^, in particular the strain Rv2571c Q335STOP (L2.2.1 from Bangladesh, ITM-500365; CT1999-00168), its *rv2571c* WT relative (L2.2.1 from South Korea, ITM-500520; CT2004-01198), as well as the strain Rv2571c V243fs (L4.1.2.1 from Nepal, ITM-500591; CT2004-01677) and its WT relative (L4.1.2.1 from Peru, ITM-500617; CT2004-02612). All strains from the BCCM collections used harbor wild-type sequences for the gene *pncA* and *panD* confirmed by sanger sequencing. Strain information and genome sequences are available at https://bccm.belspo.be. *Mycobacterium smegmatis* (Msmeg) mc^2^155 was used to express the *rv2571c* gene. Cloning and amplification of plasmids used was done using *Escherichia coli* TOP10 competent cells. A list of the plasmids, primers and compounds used in this study can be found in **Tables S4** and **S5.**

**Mycobacterial cultures**

Mtb cultures were grown at 37°C in standard Middlebrook 7H9 media containing 0.2% glycerol (~ 22mM), tyloxapol 0.05% supplemented with + homemade 10% ADN supplement (50 g/L bovine serum albumin (BSA), 20g/L dextrose, and 8.5 g/L NaCl) or on 7H10 agar media supplemented with 0.5% glycerol and 10% OADC supplement (Becton Dickinson, #212240). 4g/L of activated charcoal (Sigma Aldrich, #C9157-500G) were included into the 7H10 agar media to limit any possible carryover effects of antibiotics for the enumeration of Mtb CFUs during drug treatment from *in vitro*, ex vivo and *in vivo* experiments. 7H9 media with tyloxapol 0.05%, 5 g/L of fatty acid-free (FAF)-BSA (Sigma Aldrich, #03117057001) and 0.85 g/L NaCl were used to grow Mtb with the lipids OA, PA and CHO as main carbon source as previously described^40^. For media supplemented with OA, a stock of 1% (40 mM) oleic acid (1% [volume/volume] OA, 0.05N NaOH) was prepared in distilled water and added to cultures at a final concentration of 200 μM every two to three days. 100 mM stocks of PA and CHO were prepared in tyloxapol:ethanol (1:1), warmed to 80 °C and added at every two to three days at 200 μM (PA) or 100 μM (CHO). For the preparation of High-BSA-Oleate (HBO) media, 7H9 media with tyloxapol 0.05%, 50 g/L of FAF-BSA, 0.85 g/L NaCl and 3mM OA were used at the indicated pH (no glycerol and dextrose added). Apart from “standard” 7H9 media, the buffering agent MES (Sigma-Aldrich, #M3671) was added to acidified media (pH≤ 6.5) at a final concentration of 100 mM and the buffering agent MOPS (Sigma-Aldrich, #M1254) was added to media (pH > 6.5) at a final concentration of 100 mM. Media pH was adjusted by addition of hydrogen chloride and sodium hydroxide using a pH meter and media were filter sterilized before used. Antibiotics were added to cultures when required at the following concentrations: hyg 50 μg/mL, kan 50 μg/mL and zeo 25 μg/mL. For targeted induction of CRISPRi-based *rv2571c* silencing in Mtb, ATc was added to a final concentration of 500 ng/ml. Mtb broth cultures were maintained in standing tissue culture flasks in a humidified incubator at 37°C with 5% CO_2_. Msmeg cultures were grown similarly to Mtb at the exception that the 7H9 broth media was not supplemented with ADN.

**Generation of Mtb mutants and genetic complementation**

The mutant strains ∆*rv2571c*, ∆*hupB* and ∆*blaR*, were constructed in Mtb H37Rv by recombineering as previously described^107^. Knockout cassettes were synthesized and cloned by Genescript in a PUC57 plasmid backbone. Knockout cassettes consisted in an hygromycin cassette flanked by 500 bp homology arms corresponding to the regions immediately upstream and downstream of the target gene. For the recombineering procedure, Mtb harboring the pNitET-SacB-kan plasmid was grown to an OD_580_ of 0.6–0.8. Recombinase expression was induced by adding 1 µM isovaleronitrile for 8 hours. Electrocompetent cells were then prepared by 3 consecutive washes in 10% glycerol and an overnight glycine incubation step. A linear DNA substrate was generated by digestion of the cassettes with the restriction enzyme pmeI followed by purification on PCR purification columns (Qiagen, #28106). The cells were transformed with 500 ng of the digested substrate. To ensure the expression of flanking genes remains unaltered, 40-60bp nucleotides were preserved at the start and/or at the end of some of the deleted genes Recombinants were selected on 7H10 agar plates containing hygromycin and checked for deletion by PCR. Complemented strains Δ*hupB*::*hupB* and Δ*blaR*::*blaR* were constructed by expressing the genes *hupB* and *blaR* under their native promoter in the corresponding mutants. The mutant ∆*rv2571c* was complemented by expressing the gene *rv2571c* either under its native promoter (∆*rv2571c*::*rv2571c*) or under an *hsp60* promoter (∆*rv2571c*::*rv2571c*-OE). Native promoters for *hupB*, *blaR*, *pncA*, and *rv2571c* were defined as DNA fragments extending 280 to 500 bp upstream of the ATG start codon.

Complementation plasmids were generated using Gateway cloning technology (Life Technologies) and carried either kanamycin or zeocin resistance cassettes. These integrative vectors targeted the attL5 site (for *hupB*, *blaR*, and *pncA*) or the Tweety integration site (for *rv2571c*). The mutant ∆*rv2571c* and complemented (∆*rv2571c*::*rv2571c* and (∆*rv2571c*::*rv2571c*-OE) strains were verified by whole-genome sequencing (WGS) to confirm the desired genetic modifications and the absence of spontaneous mutations in phthiocerol dimycocerosate (PDIM) synthesis genes. Seven Rv2571c variants (**Table S3**) (Ala72fs, Arg149Cys, Phe163Ser, Asp200fs, Ala212Pro, Val243fs, and Cys337Tyr) were constructed using a site-directed mutagenesis kit (NEB, E0554S) with plasmid pGMCZt-*hsp60*-*rv2571c* as the template. Each variant was expressed in the ∆*rv2571c* background under the control of an hsp60 promoter using a zeocin-resistant Tweety-integrative plasmid. Primers and plasmids used for mutagenesis are listed in **Table S4**.

**Expression of *rv2571c* in *Mycobacterium smegmatis***

Expression of the Mtb Rv2571c in Msmeg was accomplished by electroporating the integrative plasmid pGMCK-T10M-P606-SD-*rv2571c*-HIS-PPX-GFP.This vector enables anhydrotetracycline (ATc)-inducible expression of Rv2571c as a C-terminal GFP fusion, incorporating a 10-times histidine tag and a PreScission protease cleavage site (LEVLFQ/GP) to allow for the optional removal of the GFP moiety. The expression system utilizes the TetR regulator (T10M) and the ATc-regulated promoter P606, along with a consensus Shine-Dalgarno (SD) sequence (AGGAGGTATCTCC) to maximize translation^108^. Transformant bacteria were grown to mid-log phase in the presence of kanamycin and cultured on complete 7H10 agar plates containing or not ATc (500 ng/mL). Induction of the Rv2571c-GFP fusion led to the formation of a white precipitate and media acidification. This phenotype was also observed when expressing Rv2571c alone (untagged and non-fused), confirming that the pH change was due to Rv2571c function and that the GFP fusion remains functional. To compare wild-type and mutant protein behavior, the Rv2571c-R149C variant was expressed as a C-terminal GFP fusion using an identical system (**Fig.S. 8E**). Where indicated, chlorophenol red (CPR) was added to the 7H10 plates at a final concentration of 0.004% (w/v) as a pH indicator. The list of plasmids used to express Rv2571c in Msmeg are listed in **Table S4**.

**CRISPRi chemical-genetic screening and library preparation for Illumina sequencing**

As previously described^53^, chemical-genetic screens were initiated by thawing 1 ml single-cell suspension aliquot (1 OD_580_ unit per ml) of an Mtb CRISPRi library (RLC12; Addgene 163954) and inoculating the aliquot into 29 ml 7H9 supplemented with OADC, tyloxapol 0.05% and kanamycin (10 μg/ml) in a vented tissue T-75 culture flask. The starting OD_580_ of the culture was approximately 0.1. The culture was then expanded to OD_580_ = 1.5 (~ 7 days) and use to inoculate 19 ml 7H9 supplemented with OADC, tyloxapol 0.05% and kanamycin (10 μg/ml) and ATc (100 ng/ml final concentration) to initiate target pre-depletion for 5 days. The pre-depleted culture was used to inoculate screening cultures at a starting OD_580_ of 0.05. The screen was performed in 25 ml 7H9-OA medium at pH 5 within T-75 culture flasks. Cultures were supplemented with 100 ng/mL ATc, 10 µg/ml kanamycin, and either the test compound PZA at 50 µg/mL or an equivalent volume of DMSO as a vehicle control. These cultures were grown for 14 days at 37°C with 5% CO₂, with fresh ATc replenished (100 ng/mL) at day 7 and OA replenished every 2-3 days at 200µM final concentration. After the 14-day challenge period, the cultures were harvested, washed with PBS-tyloxapol three times, and resuspended in standard 7H9 medium (near neutral pH) without any drug to allow for hit enrichment and phenotypic recovery before final collection. Genomic DNA was isolated from bacterial pellets using the CTAB‐lysozyme method as previously described^53^. The concentration of isolated genomic DNA was quantified using the DeNovix double-stranded DNA high sensitivity assay (KIT-DSDNA-HIGH-2; DS-11 series spectrophotometer/fluorometer). Next, the sgRNA-encoding region was amplified from 500 ng genomic DNA with 17 cycles of PCR using NEBNext Ultra II Q5 master mix (NEB M0544L) as previously described^53,54^. Each PCR reaction contained a pool of forward primers (0.5 μM final concentration) and a unique indexed reverse primer (0.5 μM final concentration) as previously described^53^. The resulting ~230 bp amplicon was purified using AMPure XP beads (Beckman–Coulter A63882) using one-sided selection (1.2×). Eluted amplicons were quantified with a Qubit 2.0 fluorometer (Invitrogen), and amplicon size and purity were quality controlled by visualization on an Agilent 2100 bioanalyzer (high sensitivity chip; Agilent Technologies 5067–4626). The purified amplicons were multiplexed into 10 nM pools for sequencing. To enhance sequence diversity, a 2.5–5% PhiX control spike-in (Illumina, FC-110-3001) was added. Sequencing was performed on an Illumina HiSeqX platform using single-read runs (1 × 85 cycles).

**CRISPRi data analysis and hit calling**

Sequencing counts were analyzed as previously described^53,54^. Counts were normalized for sequencing depth and an sgRNA limit of detection (LOD) cut-off was set at 100 counts in the DMSO condition. Only sgRNAs that made the LOD cut-off (i.e. counts > 100) were analyzed further. Replicate screens were quality controlled to ensure that the Pearson correlation was >0.95 for both the non-targeting sgRNA sets and essential-gene targeting sgRNA3 sets between each replicate screen. sgRNA counts were analyzed using MAGeCK (version 0.5.9.2)^53,109^ in python (version 2.7.16) comparing each drug treatment condition to the matched vehicle control (DMSO) sample. Gene-level log2 fold change (L2FC) was calculated using the ‘alphamedian’ approach specified with the ‘gene-lfc method' parameter, which estimates the gene-level L2FC as the median of sgRNAs that are ranked above the default cut off in the Robust Rank Aggregation used by MAGeCK. Negative control sgRNAs were used to calculate the null distribution and to normalize counts using the ‘– control-sgrna' and ‘–normalization control’ parameters, respectively. Unless otherwise specified, a gene was determined to be a hit in a given condition if it had a false discovery rate (FDR) < 0.01 and a log2 fold change (L2FC) < -1 in negative selection or L2FC > 1 in positive selection.

**CRISPRi-mediated *rv2571c* knockdown**

An individual CRISPRi plasmid to knockdown *rv2571c* was cloned as previously described^110^ using the Addgene plasmid 166886. Briefly, the CRISPRi plasmid backbone was digested with BsmBI (NEB R0739L) and gel purified. A sgRNA was designed to target the non-template strand of the *rv2571c* gene (targeted sequence: GACCACCGGGTCGGGCGGGAAGA) using the sgRNA design tool https://pebble.rockefeller.edu/#tools. Two complementary oligonucleotides with appropriate sticky end overhangs were annealed and ligated (T4 ligase, NEB #M0202M) into the BsmBI-digested plasmid backbone. Successful cloning was confirmed by Sanger sequencing. The resulting CRISPRi plasmid plRL58-sg*rv2571c* and non-targeting control plasmid containing a 22 mer single guide RNA not targeting Mtb genome (GCATCCGGAGCCCGTCCGTTAA) were then introduced into Mtb by electroporation of 100 ng of plasmids. sgRNA sequences and plasmids can be found in **Table S4** and S**6**.

**Antibacterial activity measurements**

All chemical compounds were dissolved in DMSO (Sigma Aldrich, #D4540), and dispensed using an HP D300e digital dispenser in a 384-well plate format. DMSO did not exceed 1% of the final culture volume and was maintained at the same concentration across all samples. For MIC assays (non-CRISPRi), cultures were growth-synchronized to late log-phase and back-diluted to an OD_580_ of 0.05 before plating. Plates were incubated standing at 37 °C with 5% CO_2_. OD_580_ was evaluated using a Molecular Devices M5 Spectramax plate reader at 12-15 days post-plating, and percent growth was calculated relative to the DMSO vehicle control for each strain. OD_580_ values from wells containing high RIF concentration (10 µg/mL) served as a “no growth” control and were subtracted to all test wells for each strain and condition tested to minimize background noise. For experiment in **Fig. S2A**, MIC were performed in 96-well plate format to allow for OA supplementation ev. 2-3 days. For CRISPRi-mediated knockdown of rv2571c, synchronized cultures of the mutant and its corresponding non-targeting sgRNA control were pre-incubated with 500 ng/mL ATc for 3 days to ensure target gene depletion prior to the experiment. IC_50_ and IC_90_ values were calculated using a nonlinear fit (Gompertz model) in GraphPad Prism. For survival assays in growth media or PBS, bacterial cultures were started at an OD_580_ of 0.05 and cultures were serially diluted in PBS containing 0.05% tyloxapol and plated on 7H10 agar supplemented with OADC and 4 g/L activated charcoal. Plates were incubated at 37 °C for 3 weeks prior to CFU enumeration.

**Culture and Metabolite Extraction in *Mycobacterium tuberculosis***

Mtb WT, ∆*rv2571c* and ∆*rv2571c*::*rv2571c*-OE strains were cultivated at 37°C until midlog phase in standard 7H9 media containing 0.05% tyloxapol (~ 50mL). The cultures were then washed twice and resuspended in phosphate buffer saline (PBS) without detergent. After measurement of the optical density to estimate bacterial number (OD_580_), approximately 5 × 10^8^ CFUs per strain were seeded onto filters and placed onto complete 7H10 plates. Plates were then incubated at 37°C for 5 days to increase biomass. The Mtb–laden filters were transferred to “swimming pools” containing 7H9 media at pH5.0 (0.5 g/L FAF-BSA, 0.085% NaCl, 200 μM OA) devoid of detergent and MES buffer. After 24h incubation at 37°C to allow Mtb to adapt to the swimming pools setup, the filters were transferred to new swimming pools of media with the same composition and pH but containing PZA at 1000 µg/mL (10x MIC_90_) or an equivalent volume of DMSO (no drug control). After 24h incubation at 37°C, Mtb–laden filters were metabolically quenched by plunging filters into a mixture of acetonitrile/methanol/H2O (40:40:20) precooled to −40°C, and metabolites were extracted by mechanical lysis with 0.1 mm zirconia beads in a Precellys tissue homogenizer for 3 min (6,500 rpm) three times under continuous cooling at or below 4 °C. Lysates were then clarified by centrifugation and collected in a new tube. After filtration across a 0.22-μm filter to ensure sterility, around 500µL of lysates and spent media from the swimming pools were collected for each condition tested and stored at -80°C. Before liquid chromatography–mass spectrometry (LC-MS), 100µL of solvent B (Acetonitrile +0.2% Formic Acid) was added to 100µL of lysates, mixed and spun down at 15,000 rpm for 8 minutes (4°C). Media samples were processed the same than lysate samples but with an additional prior step of mixing 40 μL of media with 160 μL of 50:50 Methanol:Acetonitrile followed by mixing and spinning down at 15,000 rpm for 8 minutes (4°C). 100µL supernatants of lysates and media samples were then added to capped tube to proceed with LC-MS.

**Culture and Metabolite Extraction in *Mycobacterium smegmatis***

Msmeg Wt and Msmeg*::rv2571c* were cultivated at 37°C to midlog phase in standard 7H9 media containing 0.05% tyloxapol (~ 5mL). The cultures were then washed twice and resuspended in phosphate buffer saline (PBS) without detergent. After measurement of the optical density to estimate bacterial number (OD_580_), approximately 5 × 10^8^ CFUs per strains were spread onto complete 7H10 plates containing or not ATc at 500 ng/mL (inducer of *rv2571c* gene expression). Following a 3-day incubation at 37°C, a white precipitate was observed exclusively on ATc-containing plates inoculated with Msmeg*::rv2571c*. Bacteria were removed using a cell scraper, and the underlying agar from both WT and Rv2571c-expressing plates was collected for metabolomic analysis. Agar plugs were sampled from three distinct locations per plate, flash-frozen in liquid nitrogen, and pulverized to a fine powder. The powder was resuspended at 1 mg/mL in a 4:4:2 (v/v/v) mixture of acetonitrile, methanol, and water. Proteins were precipitated by adding an equal volume of acetonitrile containing 0.2% formic acid, and the resulting supernatants were analyzed by liquid chromatography–mass spectrometry (LC–MS).

**Liquid Chromatography–Mass Spectrometry/Metabolomics analysis**

Metabolite detection was performed using liquid chromatography–mass spectrometry (LC–MS). Samples (2 μL injection volume) were separated on an Agilent 1290 Infinity LC system using a Cogent Diamond Hydride column (2.1 × 150 mm, 4 μm; Microsolv). Mobile phase A consisted of water containing 0.2% formic acid, and mobile phase B consisted of acetonitrile containing 0.2% formic acid. The flow rate was maintained at 0.4 mL min⁻¹. The 24-min normal-phase gradient was as follows: 0–2 min, 85% B; 2–5 min, 80% B; 6–7 min, 75% B; 8–9 min, 70% B; 10–11 min, 50% B; 11–14 min, 20% B; and 14–24 min, 5% B, followed by a 10-min post-run equilibration at 85% B. Mass spectrometric acquisition was carried out on an Agilent 6230 time-of-flight (TOF) mass spectrometer (Agilent Technologies) equipped with an Agilent Jet Stream electrospray ionization source, operated in extended dynamic range and negative ion mode. The capillary voltage was set to 3500 V, and the nozzle voltage was set to 1000 V. The nebulizer pressure was 35 psi, with a sheath gas flow of 11 L min⁻¹ at 350 °C. Drying gas was maintained at 250 °C at a flow rate of 10 L min⁻¹. The fragmentor voltage was set to 135 V, the skimmer to 65 V, and the octopole radio frequency voltage (Vpp) to 400 V. Data were acquired in centroid mode at 1 spectrum s⁻¹ over an m/z range of 50–1700.

**Identification of alpha-ketoglutaric acid (aKG) using the XCMS online platform**

Files generated from the LC-MS runs (d. format) were first converted into .mzML format using MSConvert before being uploaded into the XCMS online platform for untargeted metabolomics data analysis. Pairwise comparison of ATc-containing media conditioned by Msmeg::*rv2571c* versus Msmeg WT was performed in negative mode (detection of [M-H]- ions). Feature detection was conducted with a maximum tolerated m/z deviation to be 20 ppm, peaks width range of 10–60 seconds, and a signal-to-noise threshold of 10. Prefilter intensity was set at 1000 ppm. For the alignment, the bandwidth was adjusted to 10 seconds. The p-value threshold was adjusted to 0.05 and the fold change threshold to 10. Unless otherwise specified, default XCMS parameters were used. The XCMS analysis yielded 2,209 features. The features were subsequently filtered to to identify significant hits, defined as those with a ±2 log2 fold change and a q-value≤ 0.05.

**Intrabacterial pH (IBpH) measurements**

Strains expressing a pH-GFP reporter^8,39,71^ were grown to mid-log phase (OD_580_ ~ 0.5-0.8) in standard 7H9 medium. Cells were harvested by centrifugation, washed twice in PBS-tyloxapol 0.05%, and resuspended in 7mL of HBO pH5 media at a final OD_580_ of 0.1. Where indicated, the cultures were supplemented with PZA, POA, KG, Nm, Na, and Ba, or an equivalent volume of DMSO as a vehicle control. After 3 days incubation at 37°C, OD_580_ were measured to evaluate bacterial growth. The cultures were then centrifuged and resuspended in 1mL media not containing drug and the OD_580_s were measured. The OD_580_ of all suspensions were normalized to the lowest measured value by the addition of medium. Aliquots of 200µL bacterial suspension was then transferred to wells of a 96-wellfluorescence microplate. Fluorescence was measured using a Molecular Devices M5 SpectraMax plate reader, with dual excitation at 395 nm and 475 nm and emission recorded at 510 nm. IBpH was determined by calculating the 395/475 nm fluorescence ratio and interpolating these values against a calibration curve. The curve was generated using 7H9 medium with pH values ranging from 5.5 to 8.0 in 0.5-unit increments, following previously described methods.

**Alpha-ketoglutarate (αKG) quantification by fluorometric method**

The alpha-Ketoglutarate (αKG) Assay Kit (Mybiosource, MBS169589) was used to measure αKG concentrations through coupled enzymatic reactions following the protocol from the manufacturer. Briefly, αKG was transaminated to yield pyruvate, which was subsequently detected via a pyruvate-specific fluorometric probe. 50µL of samples and standards were incubated with 100µL reaction reagent in a 96-well plate and incubated for 120 minutes at 37°C protected from light. Fluorescence (Ex530nm/Em590nm) was measured using a Molecular Devices M5 Spectramax plate reader. Background-corrected fluorescence values were calculated by subtracting the average fluorescence of the zero standard from all sample readings. αKG concentrations in samples were then determined by interpolation from a standard curve.

**Structural homology search, amino acid conservation analysis and alpha-ketoglutarate (αKG) binding prediction.**

The structural predictions of Rv2571c were performed by AlphaFold Multimer^64^ using the ColabFold notebook^111^ and by AlphaFold3^112^ using the AlphaFold Server. Structural homologs were identified using AlphaFold predictions as templates to perform searches through Foldseek^113^ and Daliserver^114^. Consurf^115^ was used to generate multiple sequences, calculate conservation scores, and map scores on to predicted 3D structure of Rv2571c. Electrostatic potential for AF2 Multimer predicted structure of Rv2571c was calculated and visualized in ChimeraX^116^. The αKG-bound state of Rv2571c was predicted using DynamicBind^117^.

**Deep mutational scanning of Rv2571c**

plRL60 (pDE43-MCZtq26) was digested with NheI (NEB, cat#R3131L) and NotI (NEB, cat# R3189L) and gel purified using QIAquick Gel Extraction Kit (Qiagen, cat# 28704). PCR inserts containing *rv2571c* native promoter-PaqCI and Pleft-mScarlet-PaqCI were amplified from plRL318 and plRL319, respectively. Inserts and digested plRL60 were assembled using Gibson Assembly (NEB, cat# E2621X) to generate a “landing pad” plasmid (plRL388) for cloning the site saturated variant library for *rv2571c* (**Table S4**).

*rv2571c* site saturation variant library was synthesized by Twist Bioscience in 96-well plate format. This library encodes protein variants replacing each position of Rv2571c with every possible amino acid, including stop and synonymous mutations. Codon selection was based on codon usage for *Mycobacterium tuberculosis* (https://www.kazusa.or.jp/codon/cgi-bin/showcodon.cgi?species=83332). Library contains the mutated *rv2571c* CDSs with a downstream 20-nucleotide degenerate barcode and is flanked by two PaqCI sites. To avoid long stretches of homopolymers and constrain barcode diversity, degenerate barcodes were made based on the following rule: NNYRNNYRNNYRNNYRNNYR, where N=G/A/C/T, R=G/A, and Y=C/T.

Variant library was resuspended and pooled. plRL388 was digested with PaqCI (NEB, cat# R0745L) and gel purified using QIAquick Gel Extraction Kit (Qiagen, cat# 28704). Library inserts and digested plRL388 were ligated using Golden Gate Assembly (NEB, cat# M1100L). Assembled library was transformed into MegaX DH10B (Invitrogen, cat# C640003) and bottlenecked for an expected coverage of 75 unique barcodes per variant. Transformed cells were cultured on large LB agar (BD Difco, cat# DF0445-07-6) plates and titer plates containing 50 ug/mL Zeocin (Alfa Aesar, cat# J67140-8EQ) to calculate CFUs. Cells from large plates were harvested and plasmids were purified using HiSpeed Plasmid Maxi Kit (Qiagen, cat# 12663) to generate variant plasmid library for PacBio sequencing and subsequent transformation into Mtb.

Variant library was digested with BlpI (NEB, cat# R0585L) and PmeI (NEB, cat# R0560L) to generate linear fragments containing *rv2571c* variant CDSs and degenerate barcodes. Fragments were gel purified using QIAquick Gel Extraction Kit (Qiagen, cat# 28704) and sequenced using PacBio sequencing to associate each variant to its unique barcode. Long-read sequences were compiled to make a variant-barcode dictionary using alignparse^118^.

Library was then transformed into Mtb and outgrown in 7H9+OADC. Transformed Mtb library was back diluted and grown in HBO pH5 media and subjected to the following treatments for 14 days: no drug (control), 50 μg/mL PZA, 100 μg/mL PZA, and 500 μg/mL PZA. After treatment, cells from three biological replicates were collected for each condition. Three biological replicates were performed. Genomic DNA was isolated from collected cells using bead beating and CTAB-lysozyme treatment^119^. The concentration of isolated genomic DNA was quantified using Nanodrop (DeNovix). The barcode region was amplified from 1 ug of genomic DNA with 15 cycles of PCR using NEBNext Ultra II Q5 master mix (NEB, cat# M0544L) as described in ^102^. Each PCR reaction contained a unique forward primer (0.5 μM final concentration) and a unique indexed reverse primer (0.5 μM final concentration). The Forward primers contain a P5 flow cell attachment sequence, a unique barcode, a standard Read1 Illumina sequencing primer binding site, and custom stagger sequences designed to ensure base diversity during Illumina sequencing. The reverse primers include a P7 flow cell attachment sequence, a standard Read2 Illumina sequencing primer binding site, and unique barcodes that facilitate sample pooling for deep sequencing.

Following PCR amplification, each ~220-bp amplicon was purified using sparQ PureMag Beads with two-sided size selection (0.80× + 0.15× = 0.95×). The eluted amplicons were quantified using a Qubit 2.0 fluorometer (Invitrogen), and their size and purity were assessed by visualization on an Agilent 2100 Bioanalyzer (high-sensitivity chip; Agilent Technologies, cat. no. 5067-4626). Next, the individual amplicons were multiplexed into 15 nM pools and sequenced on an Illumina NovaSeq X Plus sequencer according to the manufacturer's instructions (single-end sequencing: 84 cycles for read 1, plus 8 cycles for the i5 index and 8 cycles for the i7 index).

Barcode counts were aligned and compiled using dms_tools^120^. Compiled barcodes were analyzed using a negative binomial regression pipeline^66^ comparing treated cells versus untreated cells. Output functional scores were visualized as a heatmap for each condition.

To identify mutations with relative functional scores that are substantially different from the median, a modified Z-score (The ASQC Basic References in Quality Control: Statistical Techniques) was calculated for each mutation. Modified Z-scores with an absolute value of greater than 3.5 were flagged as potential outliers, whereas Z-scores between -3.5 and 3.5 were considered to be deleterious, neutral, or advantageous. For each amino acid position, an intolerance score was calculated based on Zvelebil similarity scoring^121,122^ by counting differences between amino acids. Each starting amino acid is given a score of 0.1. Each difference (i.e. ‘hydrophobic’, ‘aromatic’, ‘positive’, ‘negative’, and ‘polar’) is given a score of 0.1, such that resulting substitutions that are dissimilar to the original amino acid contribute more to the intolerance score. If a mutation is considered tolerable according to our adjusted Z-score threshold, its specific score is incorporated into the initial value of 0.1. The threshold for the score was determined based on allowing methionine as the only amino acid tolerated at the start codon. For each position of Rv2571c, the scores for intolerant mutations were summed, then divided by the maximum score possible. A score of 0, therefore, indicates full tolerance in that position, whereas a score of 1.0 denotes full intolerance. Intolerance scores were visualized onto the predict structure of Rv2571c using PyMol (The PyMOL Molecular Graphics System, Version 3.0 Schrödinger, LLC.) command “loadBfacts” and ChimeraX^116^. Amino acid conservation scores and frequency of clinical mutations were also visualized on the predicted structure of Rv2571c using these tools. Plots were generated using R Studio (Posit Team) and GraphPad Prism 10.

**Measurement of *rv2571c* gene expression by RT-qPCR**

Cultures (20 mL) of Mtb WT, ∆*rv2571c*, ∆*rv2571c::rv2571c* and ∆*rv2571c::rv2571c-OE* were grown for 10 days in standard 7H9 media until OD_580_~ 1. RNA was stabilized by mixing the cultures with an equal volume of guanidinium thiocyanate buffer, followed by centrifugation at 4 °C for 10 minutes. The resulting bacterial pellets were resuspended in 1 mL TRIzol reagent and lysed by bead beating. Beads were removed by centrifugation, and the resulting supernatants were transferred to new tubes. 200 µL of chloroform were added, followed by vigorous mixing. The samples were then removed from the BSL-3 containment. RNA was purified using the RNA Clean & Concentrator-5 Kit (Zymo Research), and gDNA was removed with TURBO DNase (Ambion). Samples were subsequently repurified using the RNA Clean & Concentrator-25 Kit (Zymo Research). Complementary DNA (cDNA) was synthesized using MuLV Reverse Transcriptase (New England Biolabs) in a 25 µL reaction. A reaction lacking both the RNase inhibitor and MuLV enzyme was included to confirm the absence of genomic DNA contamination. Quantitative PCR was performed using the Light Cycler® 480 System (Roche). The housekeeping gene *sigA* (*rv2703*) served as an internal control. Primers used for RT-qPCR of *sigA* and *rv2571c* are listed in **Table S4**.

**Macrophage preparation and infection**

Bone marrow cells were isolated from female C57BL/6 mice and differentiated into BMDMs by culturing in complete DMEM supplemented with 10% heat-inactivated fetal bovine serum (FBS; Sigma), 1% HEPES (Gibco), and 30 ng/mL recombinant murine macrophage colony-stimulating factor (mCSF; Peprotech). On day 3, cultures were replenished with an additional 30 ng/mL mCSF. On day 7, macrophages were seeded into 96-well plates at a density of 8 × 10⁴ cells per well. For infections, Mtb WT, ∆*rv2571c* and ∆*rv2571c::rv2571c* were grown to early-log phase, harvested by centrifugation, washed, and resuspended in PBS-tyloxapol 0.05%. A single-cell suspension was obtained by low-speed centrifugation (800 rpm, 12 min) before dilution into DMEM. Macrophages were infected at a multiplicity of infection of 0.1. After 4 h, cells were washed twice with warm PBS to remove extracellular bacteria. Cells were then replenished with DMEM containing INH (100ng/µl), PZA (100µg/mL), or an equivalent volume of DMSO. Intracellular bacteria were enumerated on days 0, 2, and 5 by lysing macrophages with 0.01% Triton X-100 in water and plating serial dilutions of lysates on Middlebrook 7H10 agar supplemented with activated charcoal (4g/L).

**Mouse infection**

The animal experiments were performed in accordance with National Institutes of Health guidelines for housing and care of laboratory animals and according to institutional regulations after protocol review and approval by the Institutional Animal Care and Use Committee of Weill Cornell Medicine (protocol number 0601441A). Female 7- to 8-week-old C57BL/6 mice (Jackson Laboratory) were infected with ~100 CFUs of Mtb WT, ∆*rv2571c* and ∆*rv2571c::rv2571c* using an inhalation exposure system (Glas-Col). CFU burden of lungs and spleens at each time point was determined by plating dilutions of organ homogenates on 7H10 agar containing activated charcoal (4 g/L). Four mice were euthanized at each time point for each group. For the group treated with PZA, PZA at 3mg/L or 10mg/mL was added into the drinking water starting at day 21 post-infection.

**dN/dS calculations**

The ratio of nonsynonymous (dN) to synonymous (dS) substitutions was used to quantify selective pressure (dN/dS) in genes whose knockdown conferred relative PZA resistance in the CRISPRi screen. A dN/dS value <1 is indicative of negative (purifying) selection, whereas a value >1 suggests positive (diversifying) selection. dN/dS ratios were calculated using 49,481 clinical Mtb genomes from our whole-genome sequencing database compiled from publicly available datasets, as previously described^53^. The box plot in Fig. 2C summarizes the mean dN/dS ratio across all codons for each gene, while the sliding-window analysis in Fig. 2D depicts codon-level variation in dN/dS across *rv2571c*. All analyses were performed using GenomegaMap (v1.0.1).

**Phylogenetic tree building and co-occurrence of *rv2571c* variants with mutations causing antibiotic resistance**

Analysis of the association between *rv2571c* variants (SNP and indels) and drug resistance were done using the phyOverlap algorithm as previously described. In brief, we employed a phylogenetic convergence test using the phyOverlap algorithm73 (https://github.com/Nathan-dhicks/phyOverlap). FASTQ files from selected 5,413 clinical Mtb isolates genomes were aligned to H37Rv genome (NC_018143.2) using bwa (version 0.7.17-r1188). FASTQ accession numbers are provided in **Table S2**. The online resource/tool Mykrobe v0.9.012 was used to determine drug resistance. Drug resistance was claimed in clinical strains containing at least one Single-nucleotide polymorphisms (SNPs) variant known to confer resistance to either streptomycin, RIF, PZA, INH, EMB, Ofloxacin, Kanamycin or Capreomycin. Phylogenetic tree was built using FastTree (version 2.1.11 SSE3). A list of SNPs in essential genes was concatenated to build the phylogenetic tree. Indels, drug resistance-conferring SNPs, and SNPs in repetitive regions of the genome (PE/PPE genes, transposases and prophage genes) were excluded. Tree visualization was performed in iTol (https://itol.embl.de/). The Variant Proportion bar graph in Fig. 2F representing the statistical associations of *rv2571c* variants (SNP and indels) with resistance to specific drugs were assessed using Pyseer (version 1.3.10).

Associations between *rv2571c* genetic variants (SNPs and indels) and drug resistance were tested using the PySeer framework. PySeer implements linear mixed models that account for population structure and phylogenetic relatedness to estimate the effect of genetic variation on the drug-resistance phenotype. FASTQ zfiles from selected 5,413 clinical Mtb isolates genomes were aligned to H37Rv genome (NC_018143.2) using bwa (version 0.7.17-r1188). FASTQ accession numbers are provided in **Data S2**. The online resource/tool Mykrobe v0.9.012 was used to determine drug resistance. Drug resistance was claimed in clinical strains containing at least one Single-nucleotide polymorphisms (SNPs) variant known to confer resistance to either streptomycin, RIF, PZA, INH, EMB, Ofloxacin, Kanamycin or Capreomycin. Phylogenetic tree was built using FastTree (version 2.1.11 SSE3). A list of SNPs in essential genes was concatenated to build the phylogenetic tree (**Fig. 2E**). Indels, drug resistance-conferring SNPs, and SNPs in repetitive regions of the genome (PE/PPE genes, transposases and prophage genes) were excluded. Tree visualization was performed in iTol (https://itol.embl.de/). The Variant Proportion bar graph in **Fig. 2F** representing the statistical associations of rv2571c variants (SNP and indels) with resistance to specific drugs were assessed using Pyseer (version 1.3.10).

**Fig.S1 to S12**

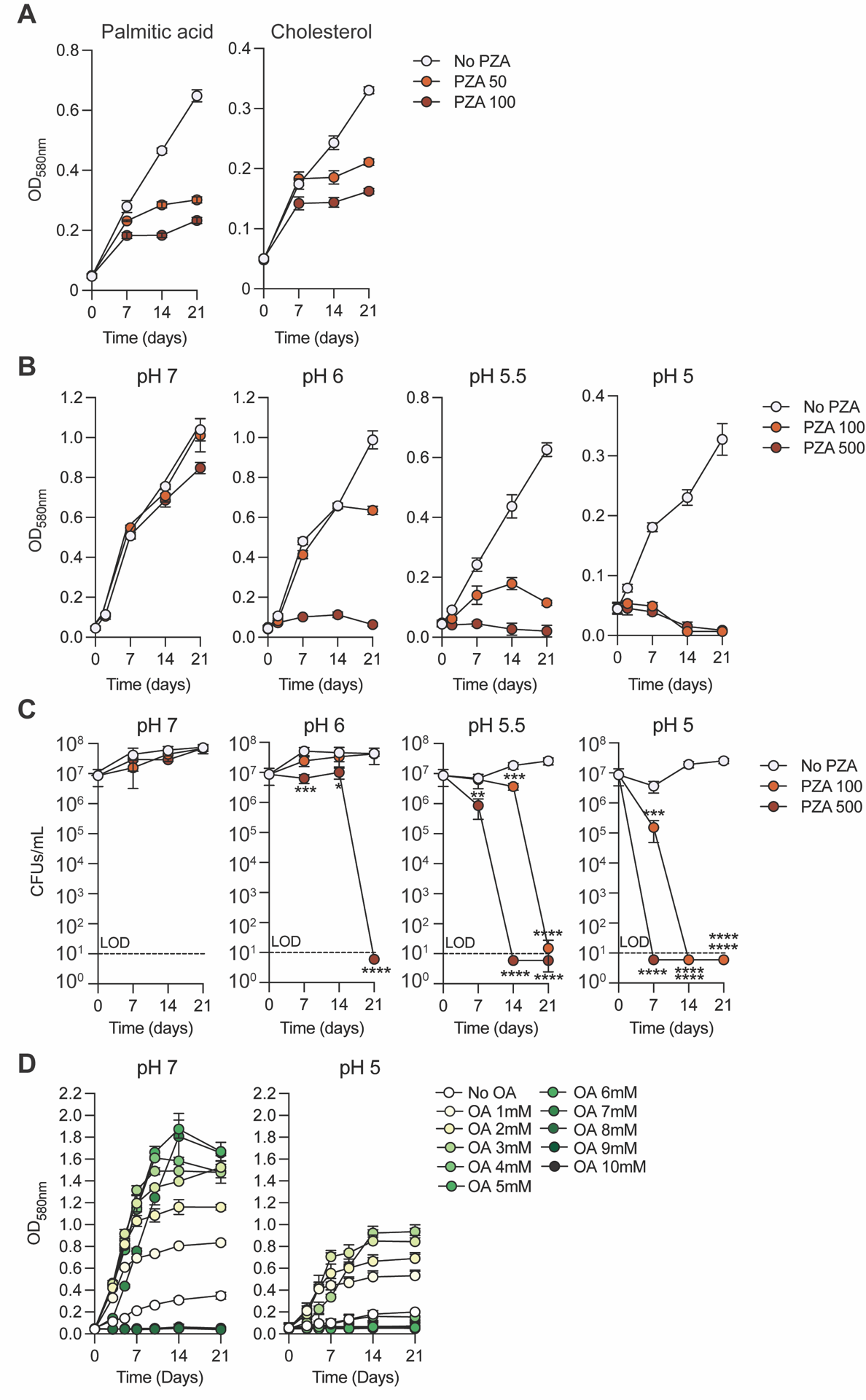

**Fig. S1. Impact of fatty acids and pH on PZA activity against Mtb**

(A) Growth of WT Mtb at pH 5.5 in media containing either Palmitic acid (PA) or Cholesterol (CHO) as the primary carbon source, treated with or without PZA (50 or 100 μg/mL).

(B-C) Growth (B) and survival (C) of WT Mtb cultured at pH 7, 6, 5.5 or 5 in the presence or absence of PZA (100 or 500 μg/mL).

(D) Growth of WT Mtb across the indicated OA concentrations in 7H9 medium supplemented with 50 g/L BSA and adjusted to pH 7 or pH 5.

In panels A–C, lipids were replenished every 2–3 days (OA and PA to 200 μM; CHO to 100 μM). Data represent means ± s.d. of three independent experiments. In panel C, statistical significance was assessed by one-way ANOVA followed by Tukey’s post-hoc test. *, adj-P < 0.05; ***, adj-P < 0.0005; ****, adj-P < 0.0001.

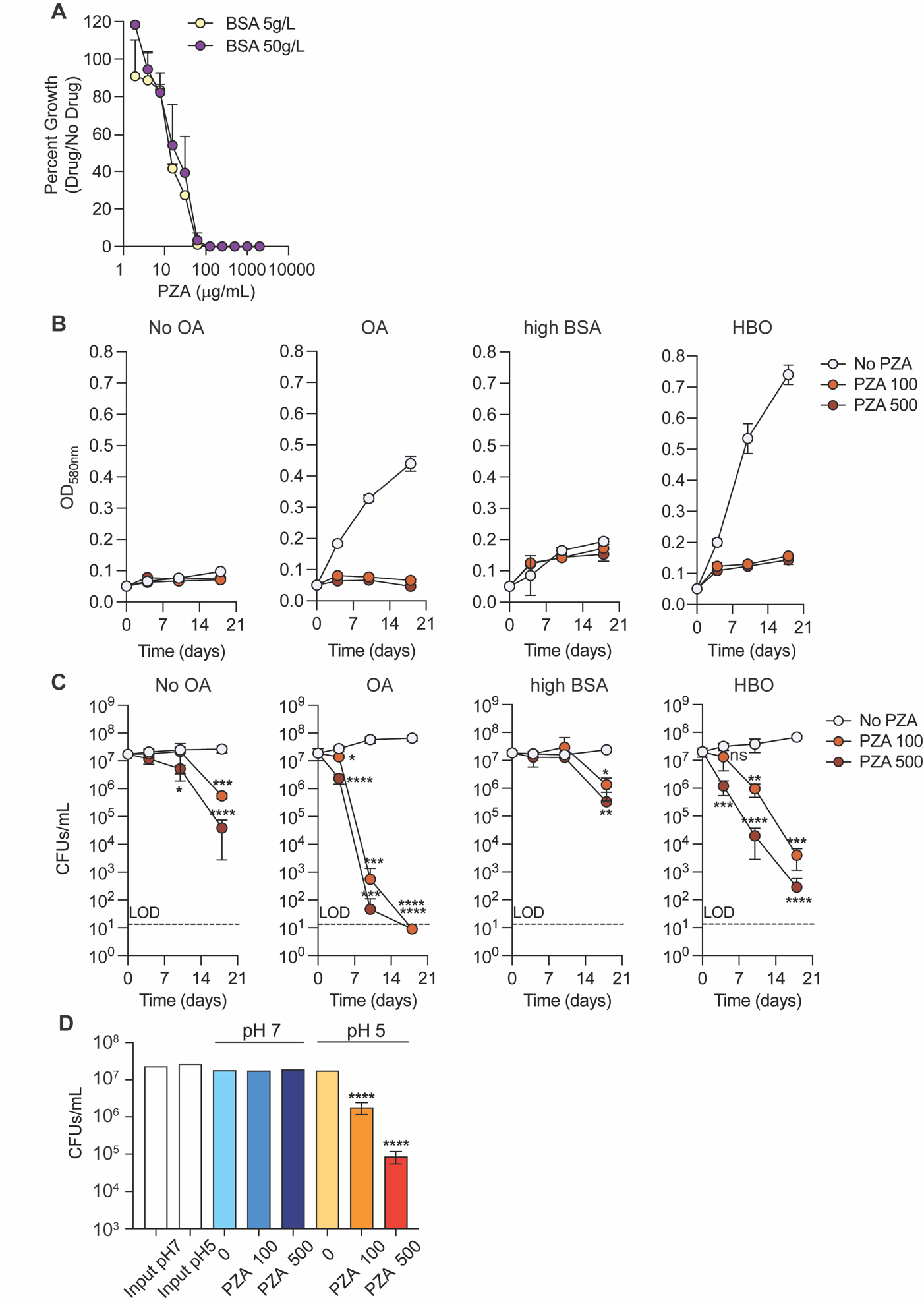

**Fig. S2. Effect of BSA concentration and medium composition on PZA activity.**

(A) PZA dose–response curves of Mtb cultured at pH 5 in the presence of standard (5 g/L) or high (50 g/L) concentrations of BSA. Data represent mean ± s.d. of an experiment performed in triplicate and representative of three independent experiments.

(B-C) Growth (B) and survival (C) of WT Mtb at pH 5 in the presence or absence of 3 mM OA, supplemented with standard or high BSA.

(D) Survival of WT Mtb in PBS buffered with 100 mM MOPS (pH 7) or MES (pH 5), treated with or without PZA (100 or 500 μg/mL).

In Panel A, B, and C, 200 μM OA was replenished every 2–3 days in “BSA 5g/L” condition (panel A) and in “OA” condition (panel B, C). In D, samples were incubated for 14 days and plated on charcoal-containing 7H10 agar for CFU enumeration. Panels B–D are means ± s.d. of three independent experiments. Statistical significance in C and D was assessed by one-way ANOVA followed by Tukey’s post-hoc test. ns, not significant; *, adj-P < 0.05; **, adj-P < 0.005; ***, adj-P < 0.0005; ****, adj-P < 0.0001.

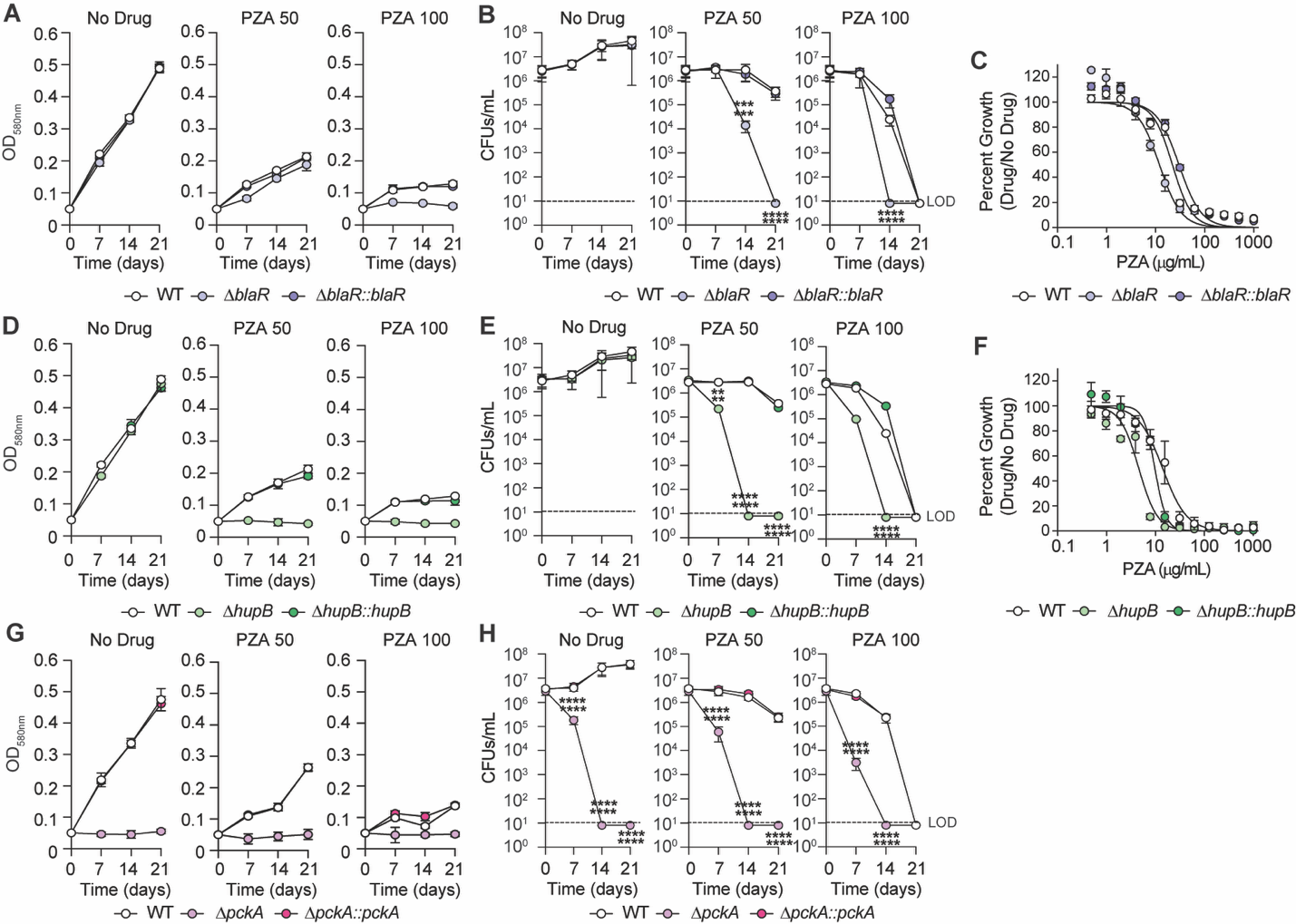

**Fig. S3. Deleting *blaR*, *hupB*, and *pckA* renders Mtb more sensitive to PZA.**

(A-B) Growth (A) and survival (B) of WT, Δ*blaR* and Δ*blaR::blaR* Mtb cultured with OA at pH 5, in the presence or absence of PZA (50 or 100 μg/mL).

(C) PZA dose–response curves of indicated Mtb strains cultured in HBO pH 5.5 medium.

(D-E) Growth (D) and survival (E) of WT, Δ*hupB* and Δ*hupB::hupB* Mtb cultured with OA at pH 5, in the presence or absence of PZA (50 or 100 μg/mL).

(F) PZA dose–response curves of indicated Mtb strains cultured in HBO pH 5.5 medium.

(G-H) Growth (G) and survival (H) of WT, Δ*pckA* and Δ*pckA::pckA* Mtb cultured with OA at pH 5, in the presence or absence of PZA (50 or 100 μg/mL).

In A, B, D, E, G, and H, 200 μM OA was replenished every 2–3 days. Data in A, B, D, E, G, and H are means ± s.d. of three independent experiments. Panels C and F are means ± s.d. of an experiment performed in triplicate and are representative of three independent experiments. In B, E, and H, statistical significance was assessed by one-way ANOVA followed by Tukey’s post-hoc test. ***, adj-P < 0.0005; ****, adj-P < 0.0001.

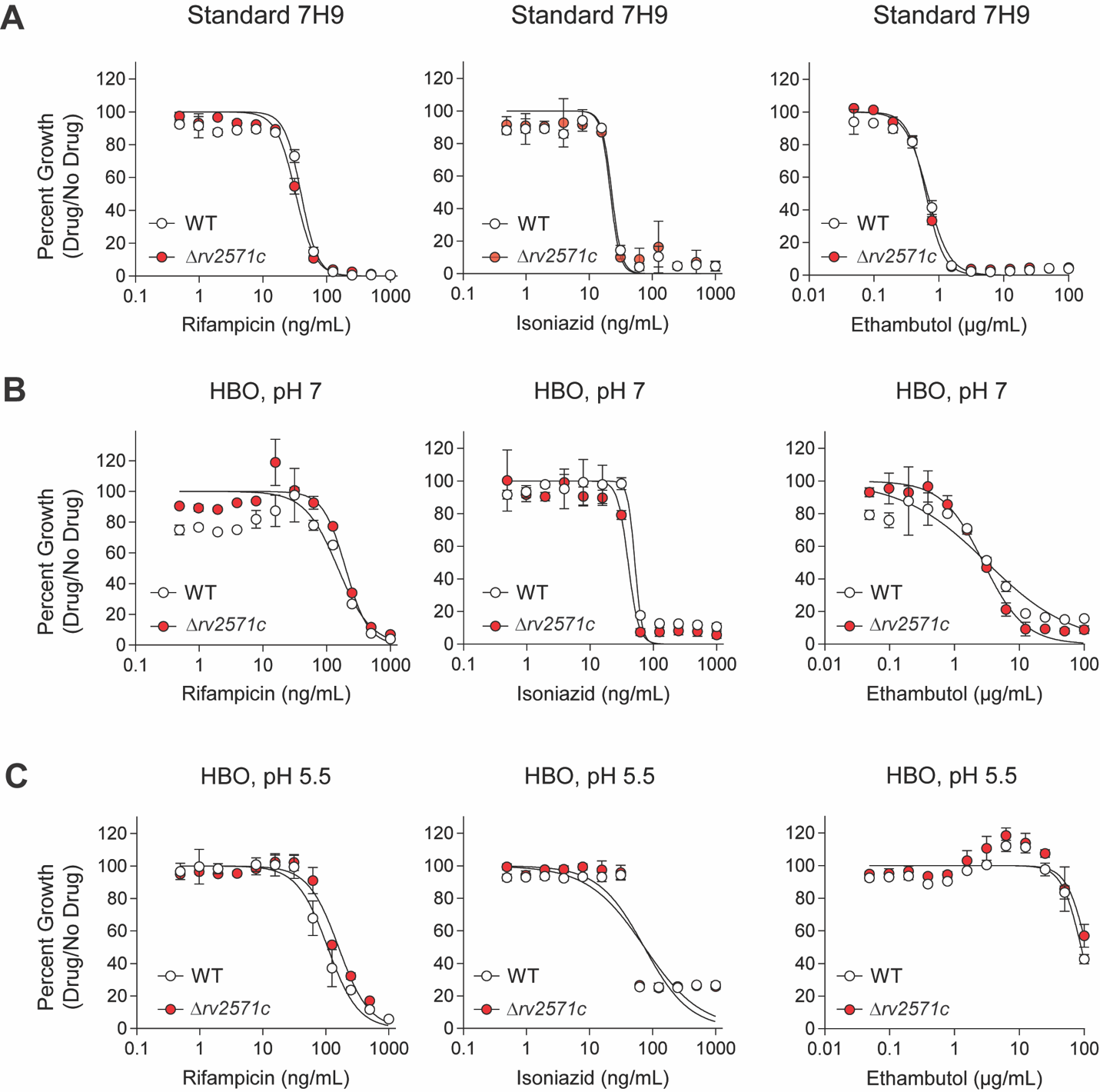

**Fig. S4. Dose–response curves of Mtb strains against first-line antibiotics across different media.**

(A–C) Dose–response curves of WT and Δ*rv2571c* Mtb treated with rifampicin, isoniazid, or ethambutol in (A) standard 7H9 medium (10% OADC, 0.2% glycerol), (B) HBO medium at pH 7, and (C) HBO medium at pH 5.5. Data are means ± s.d. of an experiment performed in triplicate and are representative of three independent experiments.

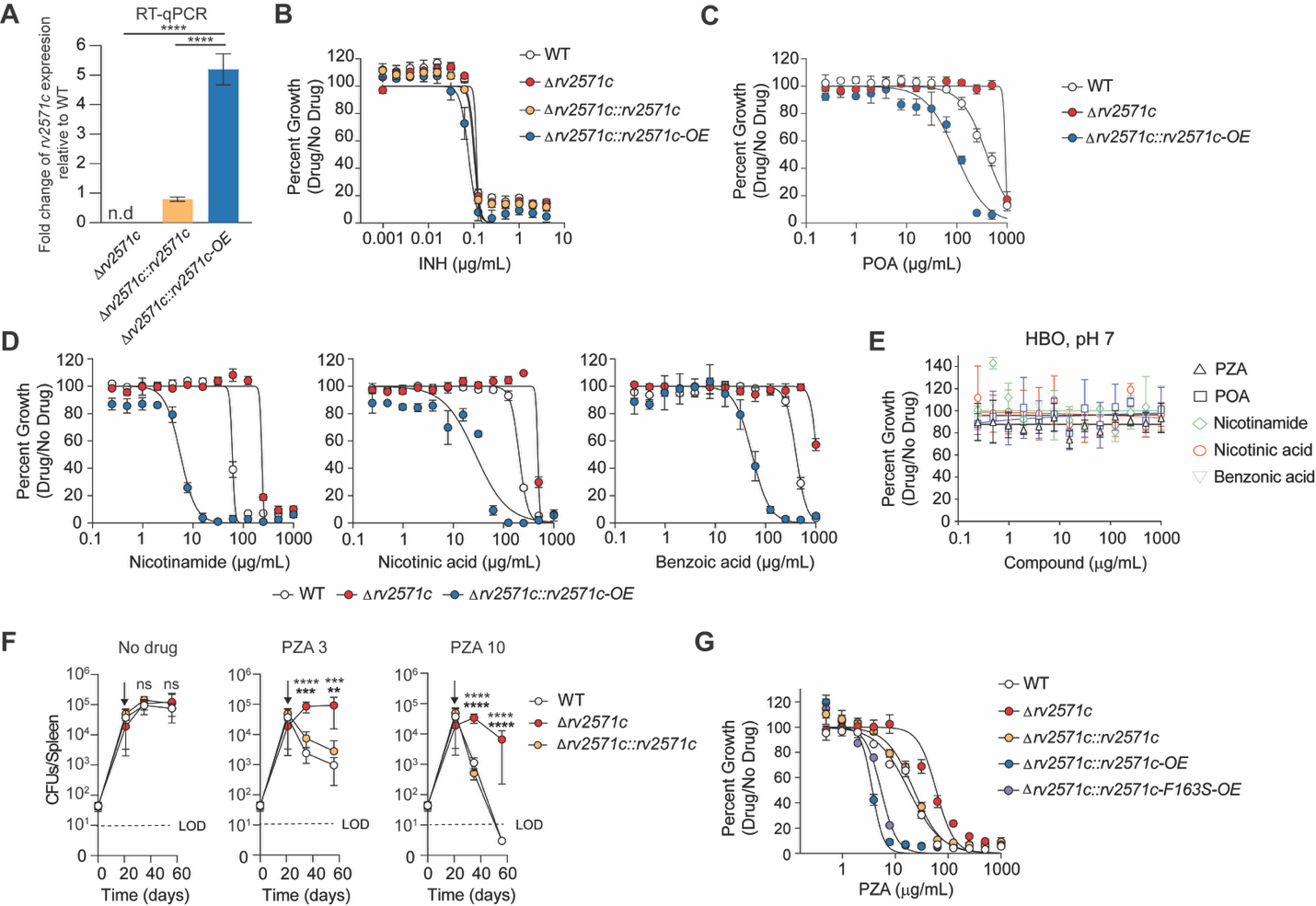

**Fig. S5. Δ*rv2571c* knockout promotes resistance to structural analogs of PZA.**

(A) Quantification of *rv2571c* mRNA levels by RT-qPCR. Indicated strains were grown for 10 days in standard 7H9 medium prior to RNA extraction.

(B–D, G) Dose–response curves of indicated Mtb strains treated with (B) INH, (C) POA, (D) nicotinamide (left), nicotinic acid (middle), or benzoic acid (right), or (G) PZA in HBO pH 5.5 medium.

(E) Dose–response curves of Mtb treated with indicated compounds in HBO pH 7 medium.

(F) Survival of indicated Mtb strains in the spleens of C57BL/6 mice. Mice were infected via aeroso and either left untreated (left) or treated with 3 mg/mL (middle) or 10 mg/mL PZA (right) in drinking water. Arrowheads indicate the start of treatment (day 21 post-infection).

(G) Dose–response curves of indicated Mtb strains treated with PZA in HBO pH 5.5 medium showing that variant *rv2571c-F163S* (fixed across lineage 3 Mtb) behaves as WT *rv2571c*.

In A–E and G, data are means ± s.d. of an experiment performed in triplicate and representative of three independent experiments. In F, data represent means ± s.d. of 4 mice per time point and are representative of two independent experiments. Dashed lines indicate the limit of detection (LOD). Statistical significance in A and F was assessed by one-way ANOVA followed by Tukey’s post-hoc test. ns, not significant; **, adj-P < 0.005; ***, adj-P < 0.0005; ****, adj-P < 0.0001.

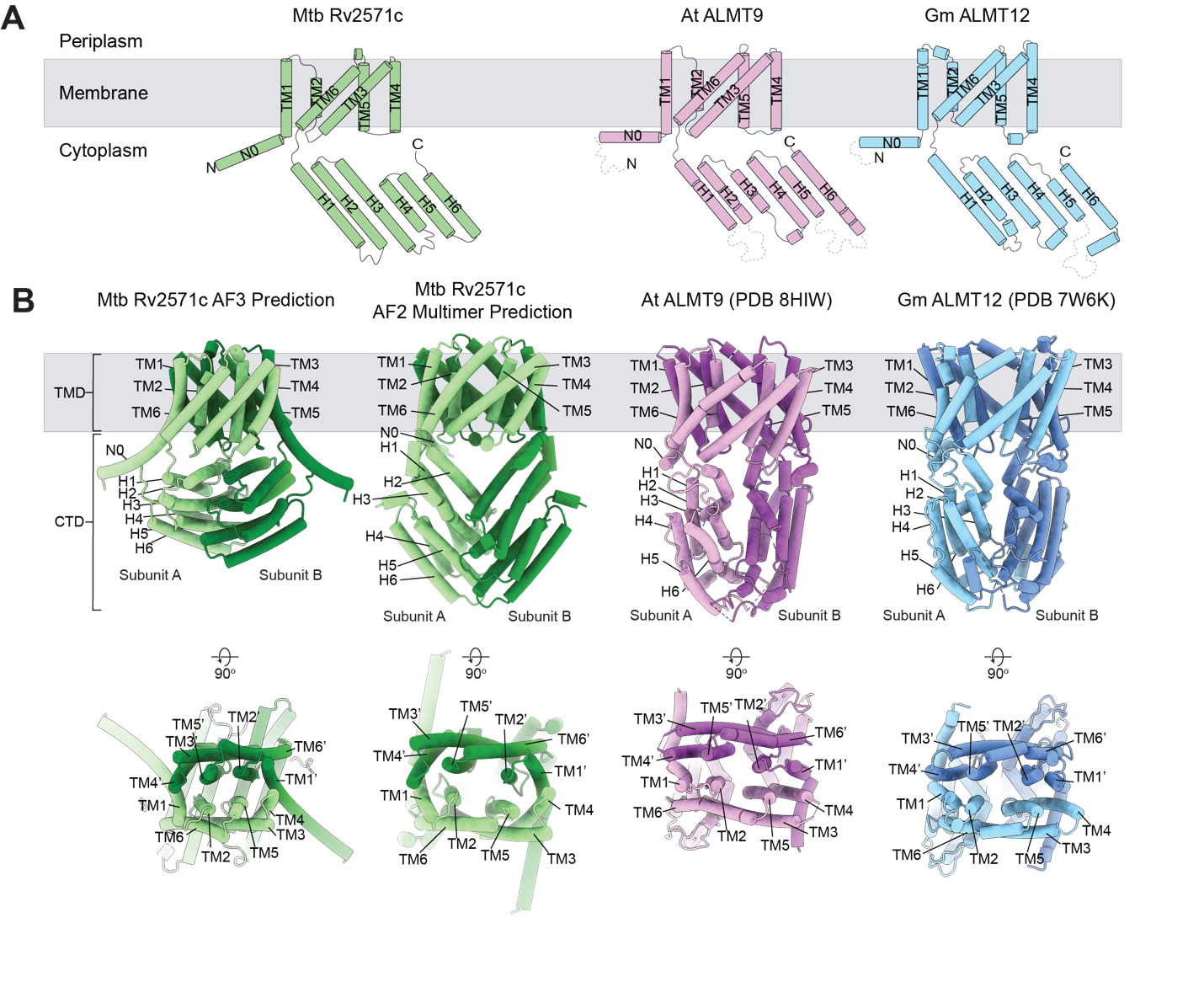

**Fig. S6. Structural conservation between Mtb Rv2571c and the ALMT protein family.**

(A) Sequence-based secondary-structure topology (wire diagram) of Mtb Rv2571c, Arabidopsis thaliana (At) ALMT9, and Glycine max (Gm) ALMT12.

(B) Predicted structures of the Mtb Rv2571c homodimer generated by AlphaFold3 and AlphaFold-Multimer compared with cryo-EM structures of At ALMT9 (PDB: 8HIW) and Gm ALMT12 (PDB: 7W6K). Structures are rendered as cylinders/stubs. The Rv2571c dimer is colored green, At ALMT9 purple, and Gm ALMT12 blue. Top row: side view of the dimers; bottom row: top-down view. The grey box indicates approximate membrane boundaries and the cytosolic region. Domains (TMD, transmembrane domain; CTD, C-terminal domain), transmembrane helices (TM), and cytosolic helices (H) are labeled.

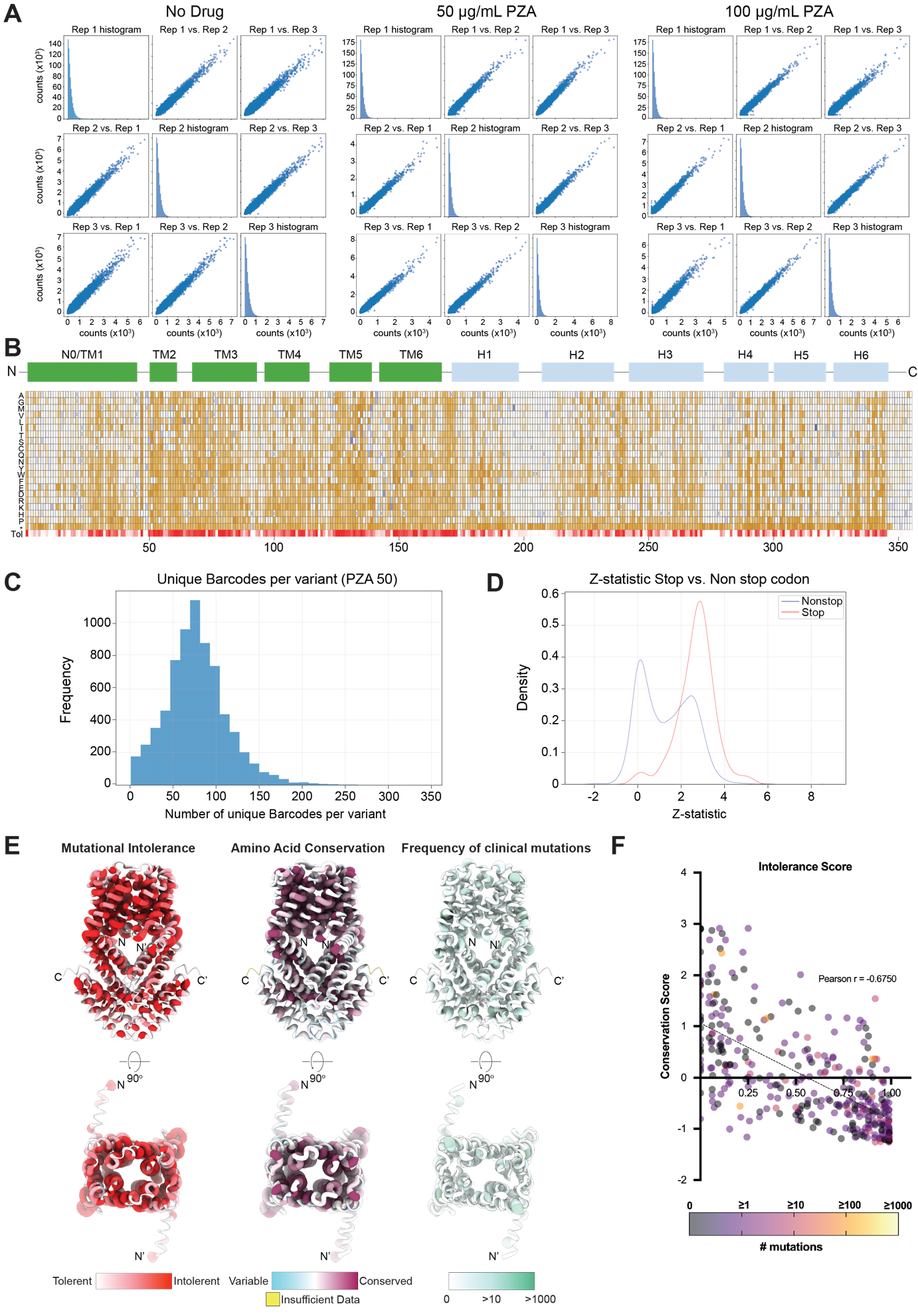

**Fig. S7. The mutational landscape of Rv2571c under PZA selection pressure.**

(A) Reproducibility of deep mutational scanning across three biological replicates (Rep 1, Rep 2, Rep 3). Scatter plots comparing barcode counts between replicates for each condition: (left) no drug, (middle) 50 µg/mL PZA, (right) 100 µg/mL PZA.

(B) Heatmaps showing functional effects (z-scores) of nearly all possible amino acid substitutions in Rv2571c under 100 µg/mL PZA selection. The secondary-structure topology of Rv2571c is shown at the top, with green boxes indicating transmembrane (TM) and light blue boxes for cytosolic (H) helices. The functional-score heatmap displays z-scores for each substitution (rows) across all positions (columns); gold indicates loss-of-function (PZA resistance), and blue indicates gain-of-function (PZA sensitivity) variants. Heatmap for mutational intolerance scores (Tol, tolerant (white) to intolerant (red)) is shown.

(C) Distribution of unique barcode counts per Rv2571c variant in the 50 µg/mL PZA condition. Each variant was represented by greater than 70 barcodes, on average.

(D) Comparison of functional z-score distributions for wildtype/missense variants versus premature STOP codons in the 50 µg/mL PZA condition.

(E) Predicted structure of the Rv2571c homodimer colored by mutational intolerance score, amino acid conservation (ConSurf), and frequency of clinical mutations, as indicated by the color legends. Structures are shown as worms; tube diameter (thickness) reflects higher scores/values.

(F) Scatter plot of mutational intolerance score versus ConSu rf conservation score. High intolerance scores (closer to 1) represent Rv2571c amino acid positions intolerant to mutation; lower conservation scores represent Rv2571c amino acid positions that are more conserved. Each data point represents one residue position in Rv2571c and is colored according to the frequency of mutations at that position observed in clinical isolates (color key below). A linear regression line (dotted) is shown.

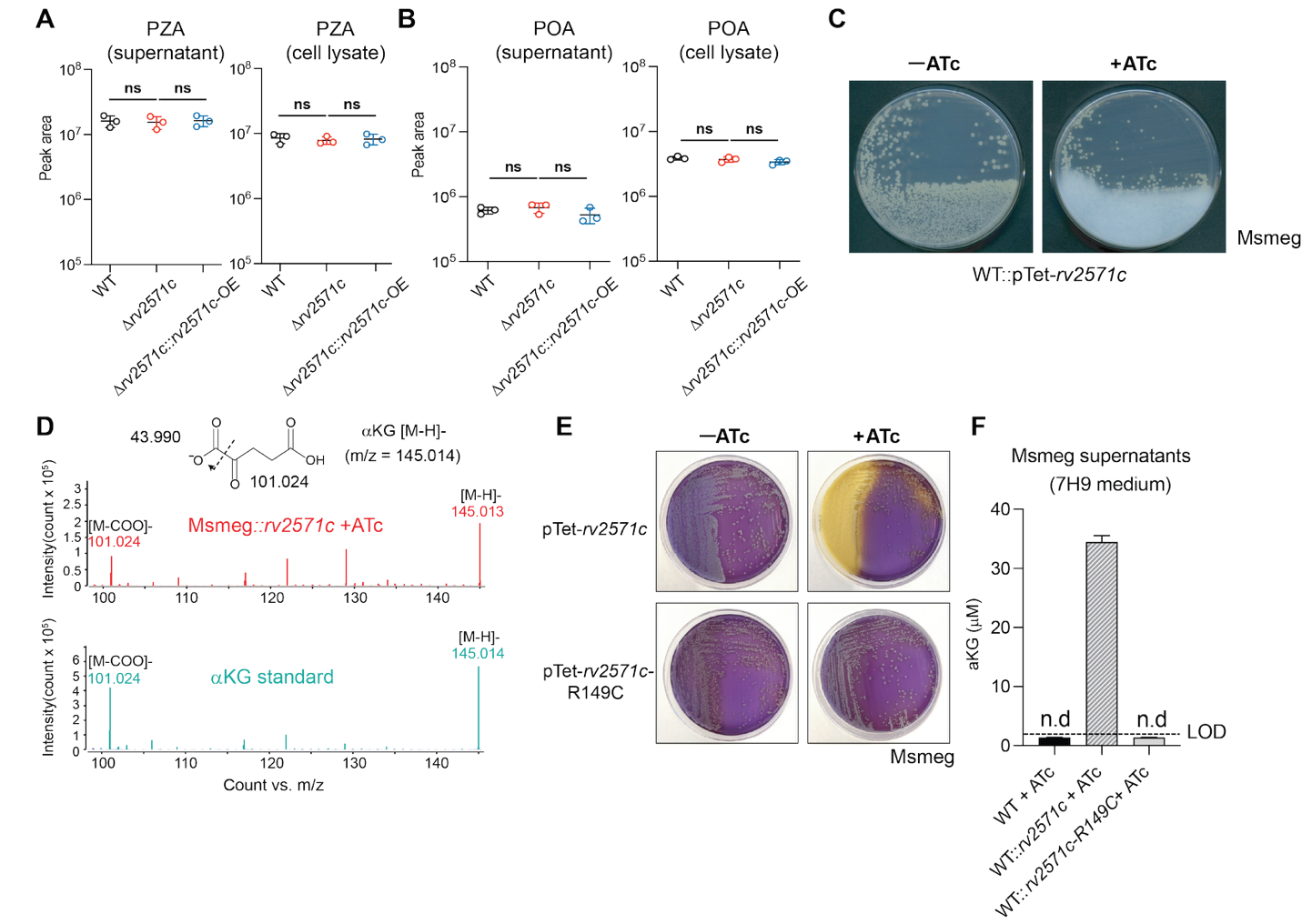
**Fig. S8. Heterologous expression of Rv2571c causes αKG secretion in *M. smegmatis*.**

(A-B) Pool size of PZA (A) and POA (B) in the indicated Mtb strains treated with PZA (10× MIC_90_) for 24 h in 7H9-OA media at pH5. Cell lysates and culture supernatants were collected and analyzed by LC-MS.

(C) Visual detection of white precipitate in solid media following ATc-induced expression of *rv2571c* in *M. smegmatis* (Msmeg::*rv2571c*).

(D) In-source fragmentation spectrum of the extract from Msmeg::*rv2571c* conditioned media (top) and a synthetic αKG standard (bottom). The feature at m/z = 101.0235 (similar to SSA) represents a fragment generated by αKG ionization under negative mode.

(E) Growth of Msmeg::*rv2571c* and Msmeg::*rv2571c*-R149C strains on chlorophenol red (CPR)-containing 7H10 plates with or without ATc (500 ng/mL).

(F) Fluorometric quantification of αKG concentration in culture supernatants of indicated Msmeg strains grown in 7H9 with ATc (500 ng/mL) for 3 days at 37°C.

Data in A, B, and F are means ± s.d. of an experiment performed in triplicate and representative of two independent experiments. Images in C and E are representative of three independent experiments. In A and B, statistical significance was assessed by one-way ANOVA followed by Tukey’s post-hoc test. ns, not significant; n.d., not detected; LOD, limit of detection.

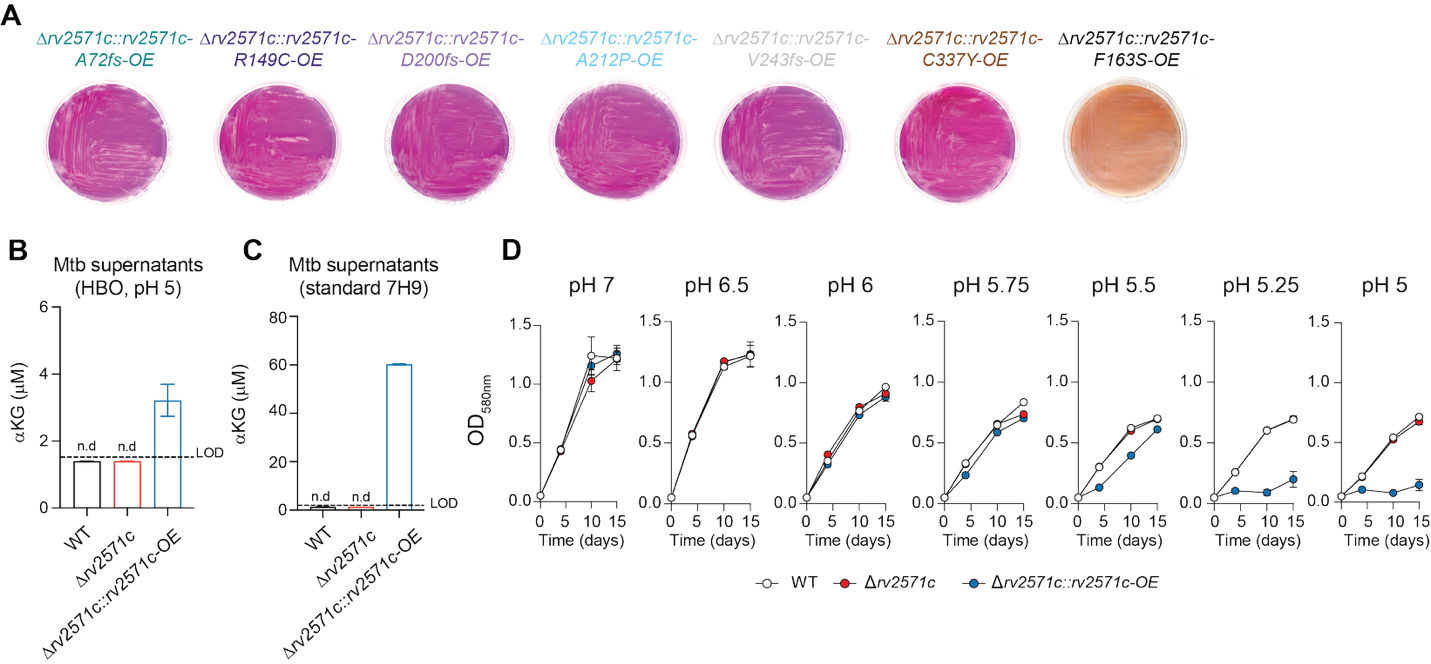

**Fig. S9. Plate acidification, increased αKG secretion and growth defect in Mtb overexpressing functional Rv2571c.**

(A) Phenotypic screening of clinically-observed *rv2571c* variants. Δ*rv2571c* Mtb strains overexpressing the indicated *rv2571c* variants were plated on 7H10 agar containing chlorophenol red (CPR) as a pH indicator.

(B-C) Quantification of αKG in supernatants of indicated Mtb strains cultured in HBO medium pH 5 (B) or standard 7H9 medium (C).

(D) Growth of indicated Mtb strains cultured in HBO medium across a range of pH values (pH 7, 6.5, 6, 5.75, 5.5, 5.25, and 5).

Images in A are representatives of three biological replicates captured after 30 days of incubation at 37 °C. Data in B and C are means ± s.d. of an experiment performed in triplicate and representative of two independent experiments. Data in D represents means ± s.d. of three independent experiments. n.d., not detected; LOD, limit of detection.

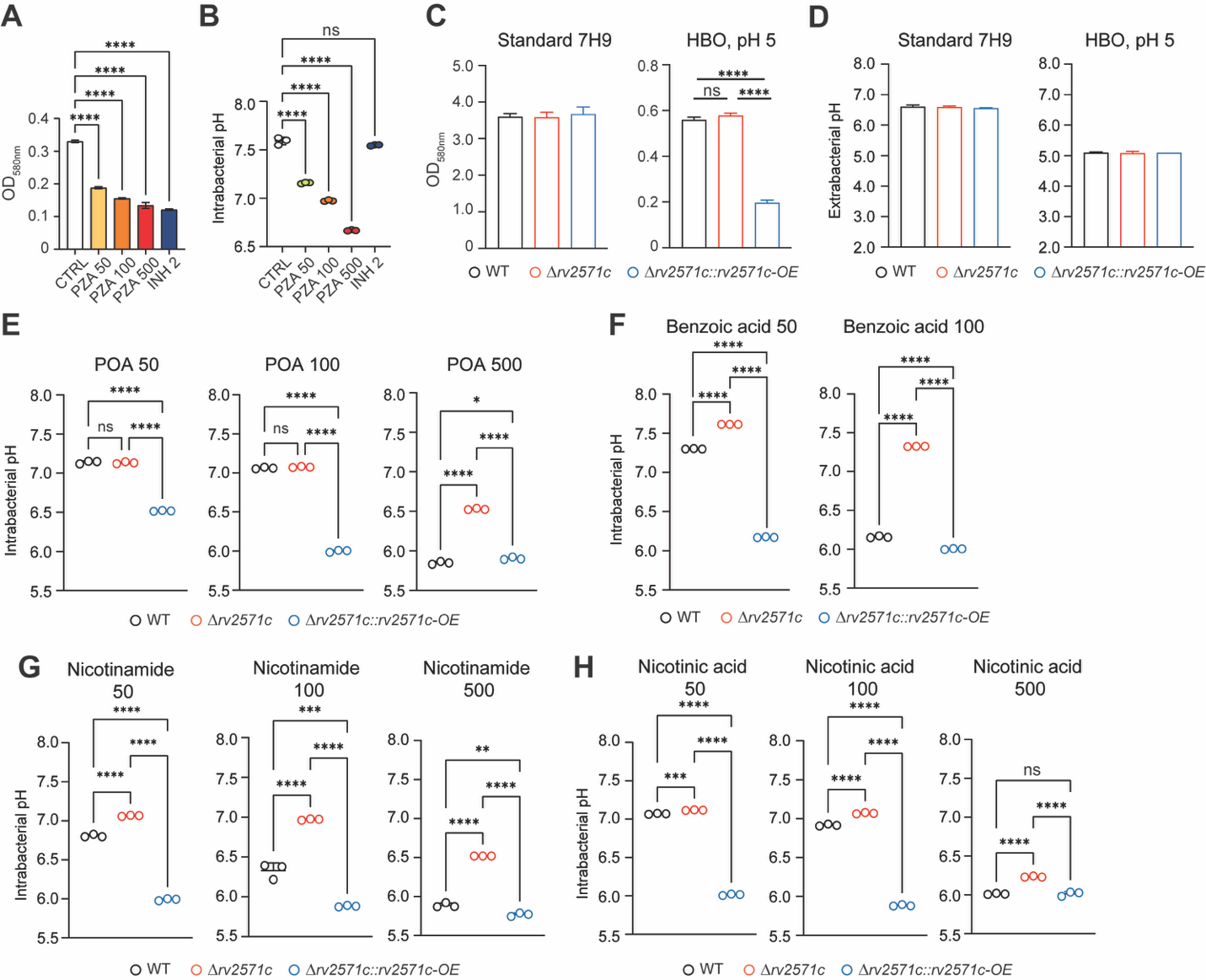

**Fig. S10. Δ*rv2571c* mitigates cytoplasmic acidification induced by PZA, POA, and their structural analogs.**

(A-B) Growth (A) and intrabacterial pH (B) of WT Mtb treated with various concentrations of PZA (50, 100, and 500 µg/mL) and INH (2 µg/mL).

(C-D) Growth (C) and extrabacterial pH (D) of the indicated Mtb strains.

(E-H) Intrabacterial pH of the indicated Mtb strains cultured in HBO medium at pH 5.5 and treated with (E) POA (50, 100, or 500 µg/mL), (F) Benzoic acid (50 or 100 µg/mL), (G) Nicotinamide (50, 100, or 500 µg/mL), or (H) Nicotinic acid (50, 100, or 500 µg/mL).

Data are mean ± s.d. from one experiment performed in triplicate, representative of three independent experiments. Significance was determined by one-way ANOVA followed by Tukey’s post-hoc test. ns, not significant; *, adj-*P* < 0.05; **, adj-*P* < 0.005; ***, adj-*P* < 0.0005; ****, adj-*P* < 0.0001.

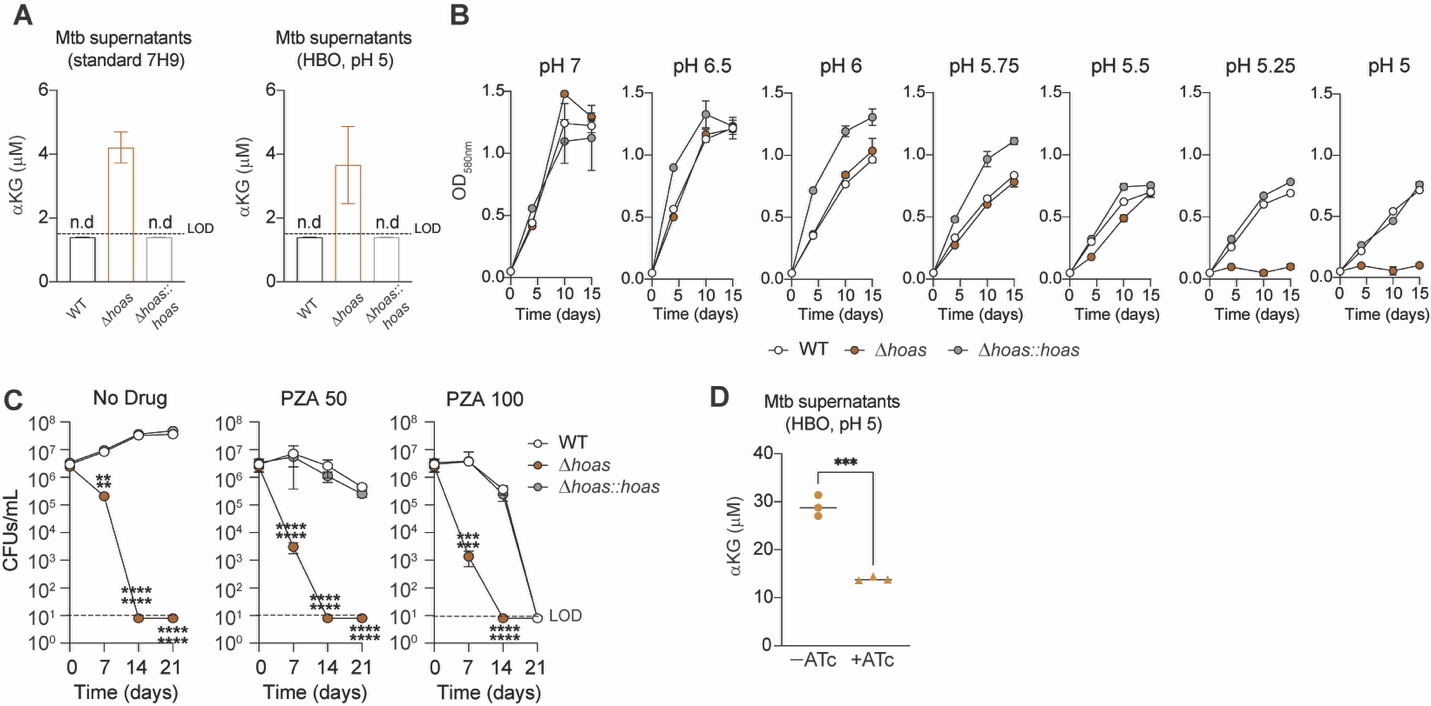

**Fig. S11. Enhanced PZA sensitivity in HOAS-deficient Mtb is associated with excessive αKG secretion.**

(A) Quantification of extracellular αKG in supernatants of indicated Mtb strains cultured in HBO medium pH 5 (left) or standard 7H9 medium (right).

(B) Growth of indicated Mtb strains cultured in HBO media across a range of pH values (pH 7, 6.5, 6, 5.75, 5.5, 5.25, and 5).

(C) Survival of indicated Mtb strains cultured with HBO medium pH 5, in the presence or absence of PZA (50 or 100 μg ml/L).

(D) Quantification of extracellular αKG in Δ*hoas* Mtb mutant harboring an sgRNA targeting *rv2571c* cultured in HBO medium pH 5.5 with or without ATc (500 ng/mL).

Data in A and D are means ± s.d. of an experiment performed in triplicate and representative of two independent experiments. Data in B and C are means ± s.d. of three independent experiments. In C, significance was determined by one-way ANOVA followed by Tukey’s post-hoc test. In D, significance was determined by Student’s t-test. n.d., not detected; LOD, limit of detection; **, adj-P < 0.005; ***, P < 0.0005; ****, adj-P < 0.0001.

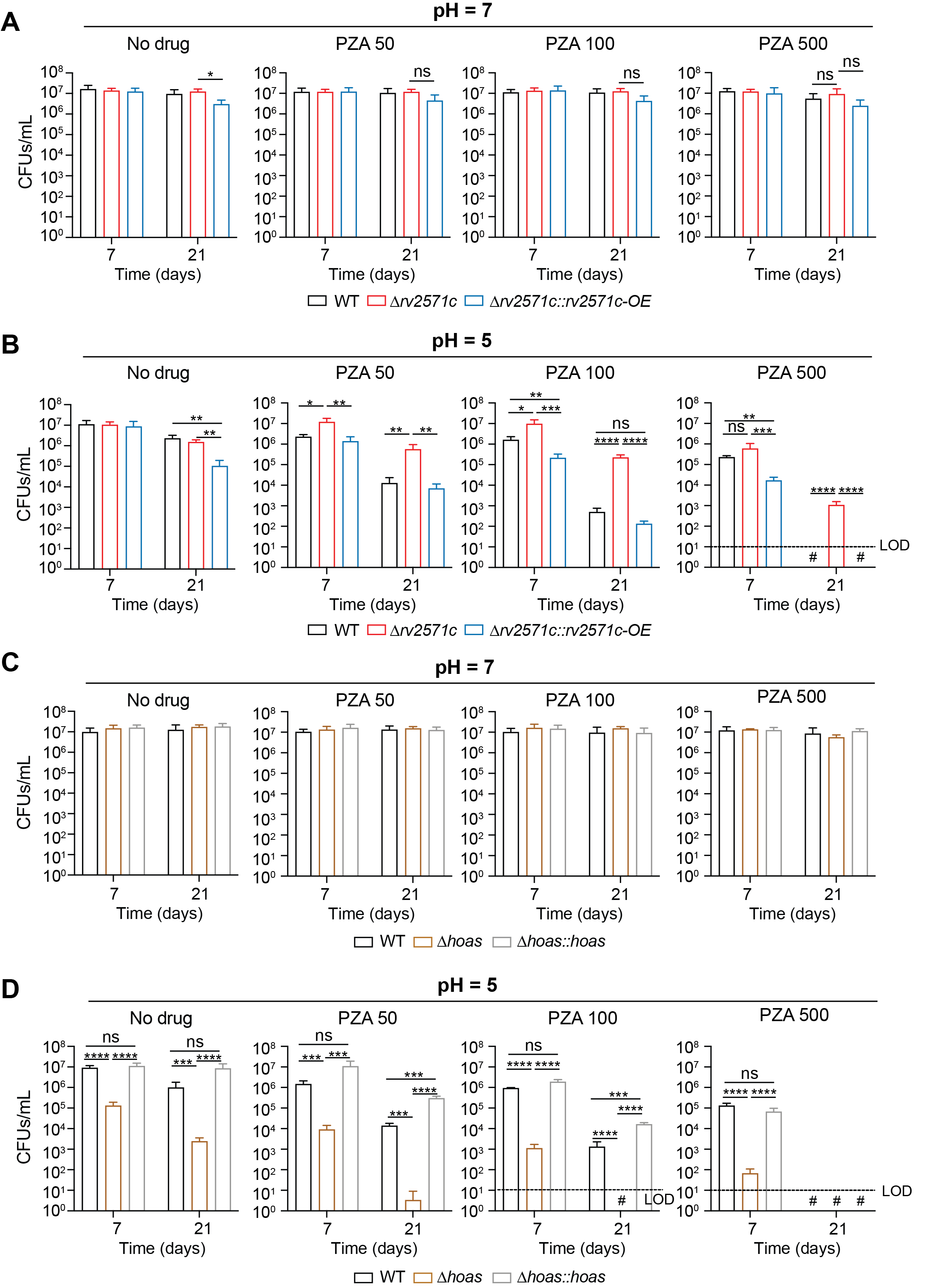

**Fig. S12. Deletion of *rv2571c* or *hoas* alters Mtb survival against PZA under non-replicative conditions.**

(A-B) Survival of WT, Δ*rv2571c,* and Δ*rv2571c*::*rv2517c-*OE Mtb strains in PBS-MOPS at pH 7 (A) or PBS-MES at pH 5 (B), treated with or without PZA (50, 100, and 500 µg/mL).

(C-D) Survival of WT, Δ*hoas,* and Δ*hoas*::*hoas* Mtb strains in PBS-MOPS at pH 7 (C) or PBS-MES at pH 5 (D), treated with or without PZA (50, 100, and 500 µg/mL).

For A–D, cultures were incubated for 3 weeks at 37 °C; bacterial suspensions were plated on charcoal-containing 7H10 agar at day 7 and day 21, and CFUs were enumerated after 3 weeks. Data are means ± s.d. of three independent experiments. Statistical significance was assessed by one-way ANOVA followed by Tukey’s post-hoc test. #, CFUs not detected; LOD, limit of detection; ns, not significant; *, adj-P < 0.05; **, adj-P < 0.005; ***, adj-P < 0.0005; ****, adj-P < 0.0001.

**Table S1 to S5**

**Table S1: List of plasmids and primers used in this work**

**Plasmids used in this work**

| **Fig. used** | **Plasmid name** | **Plasmid description** | **Resistance marker** | **Integration site** |
| --- | --- | --- | --- | --- |
| 5 | plRL58 (Addgene #166886) | Sth1 dCas9 CRISPRi plasmid optimized for use in M. tuberculosis. Sth1 dCas9 and the sgRNA are induced in the presence of ATc. This plasmid lacks the full L5 integrase and must be co-transformed plRL19. | Kanamycin | attL5 |
| 5 | plRL19 (Addgene #163634) | The L5 phage Int protein is expressed from the mycobacterial optimized promoter (MOP). This backbone is non-replicating and non-integrating in mycobacteria. | Ampicillin | / |
| 3, S7 | plJC140-Pnative-mScar-DMS-landing/ plRL388 | DMS landing pad, Integrated plasmid for complementation of rv2571c using native promoter (500bp upstream of start codon) | Zeocin | Tweety |
| 5, S10 | pGMEK_  _Ptb38_pHGFP | Expression of the pH sensitive GFP variant (pHGFP) under the strong promoter Ptb38 to measure cytoplasmic pH | Kanamycin | Episomal |
| 5 | plRL58-NT | Knock-down plasmid expressing a single guide RNA not targeting Mtb genome (Negative control). | Kanamycin | attL5 |
| 5 | plRL58-sgrv2571c | Knock-down plasmid expressing a single guide RNA to target rv2571c (targeting sequence = GACCACCGGGTCGGGCGGGAAGA; PAM = NNAGAAT) | Kanamycin | attL5 |
| 4,S8 | pGMCK_T10M_P606_SD_Rv2571c-10xHIS-PPX-GFPCter | Expression of the fusion protein *rv2571c-GFP* under the Atc inducible promoter P606 using concensus SD = AGGAGGTATCTCC. T10M allows the expression of the regulator TetR necessary for ATc regulation. Presence of 10xHIS tag and preScision protease cleaving site (PPX) sites in between *rv2571c* and *gfp* gene sequences. | Kanamycin | attL5 |
| 3,4, S8 | pGMCK_T10M_P606_SD_Rv2571c-R149C-10xHIS-PPX-GFPCter | Expression of the fusion protein rv2571cR149C-GFP under the Atc inducible promoter P606 using concensus SD = AGGAGGTATCTCC. T10M allows the expression of the regulator TetR necessary for ATc regulation. Presence of 10xHIS tag and preScision protease cleaving site (PPX) sites in between rv2571cR149C and gfp gene sequences. | Kanamycin | attL5 |
| 4,S8 | pGMCK_T10M_P606_SD_Rv2571c | Expression of the rv2571c protein under the Atc inducible promoter P606 using concensus SD = AGGAGGTATCTCC. T10M allows the expression of the regulator TetR necessary for ATc regulation. | Kanamycin | attL5 |
| S3 | pGMCK-NativeP-*blaR* | Integrated plasmid for complementation of blaR using native promoter (280bp upstream of start codon) | kanamycin | attL5 |
| S3 | pGMCK-NativeP-*hupB* | Integrated plasmid for complementation of hupB using native promoter (280bp upstream of start codon) | kanamycin | attL5 |
| S3 | pGMCK-NativeP-*pncA* | Integrated plasmid for complementation of pncA using native promoter (300bp upstream of start codon) | kanamycin | attL5 |
| 3,4,5,S5 | pGMCZt-NativeP-rv2571c (in ∆rv2571c::rv2571c) | Integrated plasmid for complementation of rv2571c using native promoter (500bp upstream of start codon) | zeocin | Tweety |
| 3,4,5,S5,S8, S9, S10 | pGMCZt-hsp60-rv2571c (in ∆rv2571c::rv2571c_OE) | Integrated plasmid for complementation of rv2571c using hsp60 promoter (overexpression) | zeocin | Tweety |
| 3,4,S9 | pGMCZt-hsp60-rv2571c-Ala72fs | Integrated plasmid for complementation of rv2571c-A72fs using hsp60 promoter (overexpression) | zeocin | Tweety |
| 3,4,S9 | pGMCZt-hsp60-rv2571c-Arg149Cys | Integrated plasmid for complementation of rv2571c-A149C using hsp60 promoter (overexpression) | zeocin | Tweety |
| 3,4,S9 | pGMCZt-hsp60-rv2571c-Phe163Ser | Integrated plasmid for complementation of rv2571c-F163S using hsp60 promoter (overexpression) | zeocin | Tweety |
| 3,4,S9 | pGMCZt-hsp60-rv2571c-Asp200fs | Integrated plasmid for complementation of rv2571c-D200fs using hsp60 promoter (overexpression) | zeocin | Tweety |
| 3,4,S9 | pGMCZt-hsp60-rv2571c-Ala212pro | Integrated plasmid for complementation of rv2571c-A212P using hsp60 promoter (overexpression) | zeocin | Tweety |
| 3,4,S9 | pGMCZt-hsp60-rv2571c-Val243fs | Integrated plasmid for complementation of rv2571c-V243fs using hsp60 promoter (overexpression) | zeocin | Tweety |
| 3,4,S9 | pGMCZt-hsp60-rv2571c-Cys337Y | Integrated plasmid for complementation of rv2571c-C337Y using hsp60 promoter (overexpression) | zeocin | Tweety |

**Primers used in this work**

| **Fig used** | **Sequence (5'-3')** | **Notes** | **Description** |
| --- | --- | --- | --- |
| 2, 3, 4, 5, S5,S8, S9, S10 | catgtttgacagcttatcatCTGGACCGCGGTGCACCA | Rv2571-KO-F1 | Rv2571 KO construct--upstream 500bp cloning |
| 2, 3, 4, 5, S5,S8, S9, S10 | tggatccactGAACGCCTATTTATTCACACTCGGGTGC | Rv2571-KO-R1 | Rv2571 KO construct--upstream 500bp cloning |
| 2, 3, 4, 5, S5,S8, S9, S10 | ataggcgttcAGTGGATCCATAACTTCGTATAATGTATG | Rv2571-KO-F2 | Rv2571 KO construct--Hyg cassette cloning |
| 2, 3, 4, 5, S5,S8, S9, S10 | tctcgccaatGGCGCGCCATAACTTCGTATAG | Rv2571-KO-R2 | Rv2571 KO construct--Hyg cassette cloning |
| 2, 3, 4, 5, S5,S8, S9, S10 | atggcgcgccATTGGCGAGAAACGCCTG | Rv2571-KO-F3 | Rv2571 KO construct--downstream 500bp cloning |
| 2, 3, 4, 5, S5,S8, S9, S10 | gtgataaactaccgcattaaAGCTATCTGCTGACAAAAAGC | Rv2571-KO-R3 | Rv2571 KO construct--downstream 500bp cloning |
| 2, S5 | CCTCTAGGGTCCCCAGCTGGCTGGACCGCGGTGCACCA | Rv2571c-endo-gibson-F1-new | complementation construct for Rv2571c KO Mtb mutant, when the newly integrated copy was expressed using native promoter (500bp upstream of Rv2571c starting codon) |
| 2, S5 | catgaccaccgggtcAGGTGGAAATAGAAGGATgctgaacacgatagc | Rv2571c-endo-gibson-R1 | complementation construct for Rv2571c KO Mtb mutant, when the newly integrated copy was expressed using native promoter (500bp upstream of Rv2571c starting codon) |
| 2, S5 | gctatcgtgttcagcATCCTTCTATTTCCACCTgacccggtggtcatg | Rv2571c-endo-gibson-F2 | complementation construct for Rv2571c KO Mtb mutant, when the newly integrated copy was expressed using native promoter (500bp upstream of Rv2571c starting codon) |
| 2, S5 | acatatccagtcactatggcTCAGGCTGACATGCCCGAC | Rv2571c-endo-gibson-R2 | complementation construct for Rv2571c KO Mtb mutant, when the newly integrated copy was expressed using native promoter (500bp upstream of Rv2571c starting codon) |
| 2, 3, 4, 5, S5,S8, S9, S10 | cctctagggtccccagctggACGGTGACCACAACGCGC | Rv2571c-hsp60-gibson-F1-new | complementation construct for Rv2571c KO Mtb mutant, when the newly integrated copy was expressed using hsp60 promoter |
| 2, 3, 4, 5, S5,S8, S9, S10 | aagcgctcatATTGCGAAGTGATTCCTCCGG | Rv2571c-hsp60-gibson-R1 | complementation construct for Rv2571c KO Mtb mutant, when the newly integrated copy was expressed using hsp60 promoter |
| 2, 3, 4, 5, S5,S8, S9, S10 | acttcgcaatATGAGCGCTTCGCTGCTAG | Rv2571c-hsp60-gibson-F2 | complementation construct for Rv2571c KO Mtb mutant, when the newly integrated copy was expressed using hsp60 promoter |
| 2, 3, 4, 5, S5,S8, S9, S10 | acatatccagtcactatggcTCAGGCTGACATGCCCGAC | Rv2571c-hsp60-gibson-R2 | complementation construct for Rv2571c KO Mtb mutant, when the newly integrated copy was expressed using hsp60 promoter |
| 3,4,S9 | GCCGATGGCACCAGCTGATCGACTGCATCAGC | 2571c-Ala212pro-F | primer used for site-directed mutagenesis to substitute alanine at position 212 to proline |
| 3,4,S9 | CAGTCGGGCGGGGCGCTG | 2571c-Ala212pro-R | primer used for site-directed mutagenesis to substitute alanine at position 212 to proline |
| 3,4,S9 | GGTTTTCGAATGTCTCTTCGACGCG | 2571c-Arg149Cys-F | primer used for site-directed mutagenesis to substitute arginine at position 149 to cysteine |
| 3,4,S9 | ACACTGCCGTTGGACGCG | 2571c-Arg149Cys-R | primer used for site-directed mutagenesis to substitute arginine at position 149 to cysteine |
| 3,4,S9 | GGATCCCACCAGCGCCCCG | 2571c-  Asp200fs-F | primer used for site-directed mutagenesis on aspartate at position 200 to cause frameshift (c.597_598insG) |
| 3,4,S9 | GCTCACCGTGTTCACCAGCTC | 2571c-  Asp200fs-R | primer used for site-directed mutagenesis on aspartate at position 200 to cause frameshift (c.597_598insG) |
| 3,4,S9 | CGTCCAAGCGTACGTCACCGATCTACAAC | 2571c-Cys337Tyr-F | primer used for site-directed mutagenesis to substitute cysteine at position 337 to tyrosine |
| 3,4,S9 | ATATCGGCGCGCACCACT | 2571c-Cys337Tyr-R | primer used for site-directed mutagenesis on cysteine at position 337 to tyrosine |
| 3,4,S9 | GGGGGGGTGCGCAGCACTG | 2571c-  Val243fs-F | primer used for site-directed mutagenesis on Valine at position 243 to cause frameshift (728delT) |
| 3,4,S9 | ATCGGCGGGGCGCTCGCC | 2571c-  Val243fs-R | primer used for site-directed mutagenesis on Valine at position 243 to cause frameshift (728delT) |
| 3,4,S9 | GGCTATCGTGTCCAGCATCCTTC | 2571c-Phe163Ser-F | primer used for site-directed mutagenesis to substitute proline at position 163 to serine |
| 3,4,S9 | AGCCCACCACCGACC | 2571c-Phe163Ser-R | primer used for site-directed mutagenesis to substitute proline at position 163 to serine |
| 3,4,S9 | GCCCAACAGATGATCGTCGGGG | 2571c-ala72fs-F | primer used for site-directed mutagenesis on Alaine at position 72 to cause frameshift  (c.213delT) |
| 3,4,S9 | CGTCGTGCGCGCAGCACG | 2571c-ala72fs-R | primer used for site-directed mutagenesis on Alaine at position 72 to cause frameshift  (c.213delT) |
| S5 | GTGATTTCGTCTGGGATGAAGA | Mtb-qPCR-sigA-F | qPCR primers used for RT-PCR assay |
| S5 | TACCTTGCCGATCTGTTTGAG | Mtb-qPCR-sigA-R | qPCR primers used for RT-PCR assay |
| S5 | TTGTGGCAGCAACGAAGT | Rv2571-qPCR-F1 | qPCR primers used for RT-PCR assay |
| S5 | CGTTCGATTACCCGTTGTAGAT | Rv2571-qPCR-R1 | qPCR primers used for RT-PCR assay |

**Table S2: List of sgRNAs used in this work**

| **Fig. used** | **sgRNA ID** | **Gene targeted** | **sgRNA targeting sequence (5'-3')** |
| --- | --- | --- | --- |
| 5 | Non-targeting control | NA | GCATCCGGAGCCCGTCCGTTAA |
| 5 | *rv2571c* sgRNA | *rv2571c* | GACCACCGGGTCGGGCGGGAAGA |

**Table S3: List of compounds and antibiotics used in this work**

| **Compound** | **Use** | **Abbreviation** | **Product number** |
| --- | --- | --- | --- |
| Pyrazinamide | Axenic culture, macrophage infection, mouse efficacy studies | PZA | Thermo Fisher, #157641000 |
| Pyrazinoic acid | Axenic culture | POA | Sigma-Aldrich, #P56100 |
| Nicotinamide | Axenic culture | Nm | Sigma-Aldrich, #72340 |
| Nicotinic acid | Axenic culture | Na | Sigma-Aldrich, #72309 |
| Benzoic acid | Axenic culture | Ba | Sigma-Aldrich, #242381 |
| Oleic acid | Axenic culture | OA | Millipore Sigma, #OX0165 |
| Palmitic acid | Axenic culture | PA | Cayman Chemical, #10006627 |
| Cholesterol | Axenic culture | CHO | Sigma-Aldrich, #C3045 |
| Sodium Butyrate | Axenic culture |  | Sigma-Aldrich, #303410 |
| Tyloxapol | Axenic culture, macrophage infection, mouse efficacy studies |  | Sigma-Aldrich, #T8761 |
| Hygromycin | Axenic culture | Hyg | Sigma-Aldrich, #10843555001 |
| Kanamycin | Axenic culture | Kan | Sigma-Aldrich, #K0254 |
| Zeocin | Axenic culture | Zeo | Thermo Fisher, #R25001 |
| Anhydrotetracycline | Axenic culture | ATc | Sigma-Aldrich, #37919 |
| Chlorophenol red | Axenic culture | CRP | Sigma-Aldrich, #199524 |
| Dimethyl sulfoxide | Axenic culture | DMSO | Sigma Aldrich, #D4540 |

**Table S4: Non-synonymous variants of Rv2571c tested in this study**

| **Variant** | **Predicted outcome** | **Nucleotide Change** | **Codon Usage** |
| --- | --- | --- | --- |
| Ala72fs | frameshift | 213delT |  |
| Arg149Cys | misense | 445C>T | cgc to tgs |
| Phe163Ser | misense | 488T>C | ttc to tcc |
| Asp200fs | frameshift | 597_598insG |  |
| Ala212pro | misense | 634G>C | gcc to ccc |
| Val243fs | frameshift | 728delT |  |
| Cys337Tyr | misense | 1010G>A | tgt to tat |

**Table S5: Influence of pH on PZA activity against Mtb in HBO medium (/: no activity)**

| **pH** | **7** | **6.5** | **6.25** | **6** | **5.75** | **5.5** | **5.25** | **5** |
| --- | --- | --- | --- | --- | --- | --- | --- | --- |
| PZA MIC_90_ (µg/mL) | / | / | / | 534 | 594 | 392 | 137 | 79 |
| PZA MIC_50_ (µg/mL) | / | / | 935 | 52 | 35 | 41 | 45 | 28 |

**Data S1 to S4**

**Data S1:** MAGeCK screen results.

**Data S2:** List of identified rv2571c mutations from clinical isolate WGS database and their co-ocurrence with known drug-confering mutations.

**Data S3:** Deep Mutational Screen (DMS) results.

**Data S4:** xcms data from supernatant expressing Rv2571c in *M.smegmatis*.
